# Hypoxia-induced ALDH7A1 expression protects colorectal cancer cells from oxidative stress and DNA damage via a HIF-1-independent mechanism

**DOI:** 10.64898/2026.08.15.729690

**Authors:** Lina Elsalem, Simon J. Allison, Maria Sadiq, Abdullahi M. Dauda, Krishna Khullar, Mark Sutherland, Steve D. Shnyder, Syed A. Khurram, Roger M. Phillips, Jan S. Moreb, Sneha Smarakan, Klaus Pors

## Abstract

Tumour hypoxia is associated with increased invasiveness, metastasis, and drug resistance; however, its impact on drug-metabolising enzymes remains poorly understood. This study investigated the effect of hypoxia on the expression of selected aldehyde dehydrogenase (ALDH) isoforms (ALDH1A1, 1A2, 1A3, 1B1, 2, 3A1, and 7A1) in colorectal cancer (CRC) cells. CRC cell lines (HT29, DLD-1, SW480, and HCT116) were cultured under normoxic and hypoxic (0.1% O₂) conditions, while HT29 and DLD-1 cells were additionally grown as multicellular spheroids (MCS). Expression of ALDH isoforms was assessed at the mRNA and protein levels. Functional studies included siRNA-mediated knockdown of ALDH1A1, ALDH3A1, and ALDH7A1, measurement of reactive oxygen species (ROS), and stable overexpression of ALDH7A1 in H1299 cells. ALDH7A1 was consistently upregulated at both transcript and protein levels in HT29 and DLD-1 cells exposed to hypoxia. Elevated ALDH7A1 expression was also observed in hypoxic regions of MCS and CRC xenografts (HT29, DLD-1, HCT116, SW620, and COLO205). Knockdown of ALDH7A1 in DLD-1 cells reduced proliferation, increased ALDH3A1 expression, and significantly elevated ROS levels, indicating a role in redox homeostasis and suggesting functional crosstalk between these isoforms. Conversely, stable overexpression of ALDH7A1 in H1299 cells markedly reduced ROS levels. Taken together, these findings identify ALDH7A1 as a hypoxia-responsive enzyme that promotes adaptation to oxidative stress and may contribute to CRC cell survival within the hypoxic tumour microenvironment.

## Introduction

Colorectal cancer (CRC) is the second leading cause of cancer-related deaths and the third most common malignancy worldwide, with an incidence of approximately 1.9 million cases globally [1–3]. Epidemiological studies reveal some association of CRC incidence with age, lifestyle, obesity, smoking and lack of physical exercise, along with a genetic predisposition [4]. Improvements in outcomes for patients with metastatic disease remain modest despite significant advances in understanding the molecular mechanisms underlying CRC pathogenesis and the introduction of targeted therapies, including monoclonal antibodies against EGFR (Cetuximab) and VEGF (Bevacizumab). The integration of targeted agents with chemotherapy and the identification of actionable biomarkers such as RAS wild-type status, BRAF V600E mutations, and HER2 amplification have enabled more precise patient stratification and improved responses in selected subgroups; however, durable and broadly effective treatment strategies are still lacking [5–7]. Several reports suggest the existence of an intricate crosstalk between the tumour microenvironment (TME) and CRC cells, inflicting profound effects on cancer progression, metastasis and drug resistance [8]. Tumour hypoxia (low pO_2_) is one of the key microenvironmental features that can affect cellular expression programs, contributing to clinical resistance in most solid tumours [9]. The major mechanism mediating adaptive responses to low O_2_ availability is changes in gene expression through the stabilisation of the oxygen-labile transcription factor, Hypoxia Inducible Factor-1 (HIF1) [9,10]. In addition, hypoxia has been reported to increase reactive oxygen species (ROS) and oxidative stress, thereby promoting tumour progression [11,12]. Several studies indicate that hypoxic cancer cells exposed to oxidative stress develop adaptive antioxidant responses or strategies to survive the hostile milieu, which may result in increased aggressiveness [12]. These antioxidant mechanisms induced by elevated ROS levels might help protect cells against radiotherapy and chemotherapeutic agents such as cisplatin, doxorubicin, and etoposide [13–15].

The human aldehyde dehydrogenase (ALDH) superfamily consists of 19 isozymes that primarily catalyse the conversion of a range of endogenous and exogenous aldehydes to their corresponding carboxylic acids, with implications for physiological functions and cellular protection [16]. In cancer, ALDH expression has been shown to influence drug sensitivity and clinical prognosis, and its functional activity has also been used to identify and isolate stem-like subpopulations [17]. Accumulating evidence supports important roles for several ALDH isoforms in CRC progression. These include ALDH1B1 as a potential diagnostic and prognostic biomarker [18], ALDH1A1 as a marker of cancer stem cells [19], ALDH1A3 as a mediator of drug resistance [20], and ALDH3A1 as a promoter of metastasis [21]. Genomic studies have further shown that members of the ALDH superfamily are upregulated in response to oxidative stress [22], enhancing cellular protection against oxidative damage induced by environmental toxins and anticancer agents [23]. Human ALDH7A1, also known as antiquitin, has been reported to protect cells against hyperosmotic stress, likely by generating betaine, an important cellular osmolyte produced by oxidation of betaine aldehyde [24]. Furthermore, ALDH7A1 has been implicated in cellular antioxidant defence mechanisms by detoxifying reactive aldehydes generated during oxidative stress. Brocker et al. demonstrated that ALDH7A1 activity confers cytoprotection under oxidative stress, in which increased lipid peroxidation (LPO) leads to the accumulation of cytotoxic aldehydes. Mechanistically, the detoxification of lipid peroxidation (LPO)-derived aldehydes by ALDH7A1 is thought to confer multiple cytoprotective benefits. By metabolising these reactive aldehydes, ALDH7A1 reduces the reliance on glutathione (GSH)-dependent detoxification pathways, thereby preserving intracellular GSH levels and enhancing cellular resistance to oxidative stress-induced damage [25].

In this study, two-and three-dimensional CRC *in vitro* models were utilised to evaluate the impact of hypoxia on ALDH isoform expression. The findings suggest that ALDH7A1 contributes to tumour cell adaptation to oxidative stress by supporting cell survival and regulating ROS levels. These observations highlight a potential role for ALDH7A1 in CRC biology and support further investigation into its utility as a biomarker and therapeutic target.

## Material and Methods

### Cell culture and chemotherapeutic agents

Human CRC cell lines DLD-1, HCT116, HT29, SW620, COLO205, and SW480, together with the human non-small cell lung carcinoma cell line H1299, were obtained from the American Type Culture Collection (ATCC, Manassas, VA, USA). DLD-1, HCT116, HT-29, and SW480 were used for *in vitro* profiling, while SW620 and COLO205 were additionally sourced for cell line-derived xenograft evaluations. The H1299/RFP and H1299/ALDH7A1 cell lines were generated as previously described [26]. All cell lines were maintained in RPMI-1640 medium (Sigma-Aldrich, UK) supplemented with 10% foetal bovine serum and 2 mM L-glutamine and cultured at 37°C in a humidified atmosphere containing 5% CO₂.

Oxaliplatin, irinotecan, and 5-fluorouracil (5-FU) were purchased from Sigma-Aldrich. For hypoxic treatment, cells were cultured in a hypoxia workstation (Whitley H35 Hypoxystation; 0.1% O₂, 5% CO₂, 95% N₂, and 95% humidity) for 24 or 48 h before RNA and protein extraction. Before hypoxic exposure, culture medium was pre-equilibrated under hypoxic conditions overnight and replaced with fresh medium immediately before treatment. Cells were harvested at approximately 75% confluency. Control cells were maintained under normoxic conditions (37°C, 5% CO₂, and 95% relative humidity) for the corresponding treatment periods.

### Quantitative polymerase chain reaction (qPCR)

Total RNA was extracted from CRC cell lines using the RNeasy Mini Kit (QIAGEN) according to the manufacturer’s instructions. cDNA was synthesised using the AffinityScript qPCR cDNA Synthesis Kit (Agilent Technologies) and diluted 1:10 before qRT-PCR. Primer sequences are listed in Appendix Table 1. qRT-PCR reactions (20 μL) contained 300 nM forward and reverse primers for *ALDH7A1*, *VEGFA*, or *β-actin*, 10 μL Precision™ 2× Master Mix, 4 μL RNase-/DNase-free water (PrimerDesign), and 5 μL diluted cDNA. *VEGFA* was used as a positive control for hypoxia and *β-actin* as the housekeeping gene. No-reverse-transcription and no-template controls were included in all assays. Reactions were performed on an Applied Biosystems 7500 Real-Time PCR System (Thermo Fisher Scientific). Relative gene expression was determined using the 2^−ΔΔCt^ method and normalised to *β-actin*.

**Table 1:**
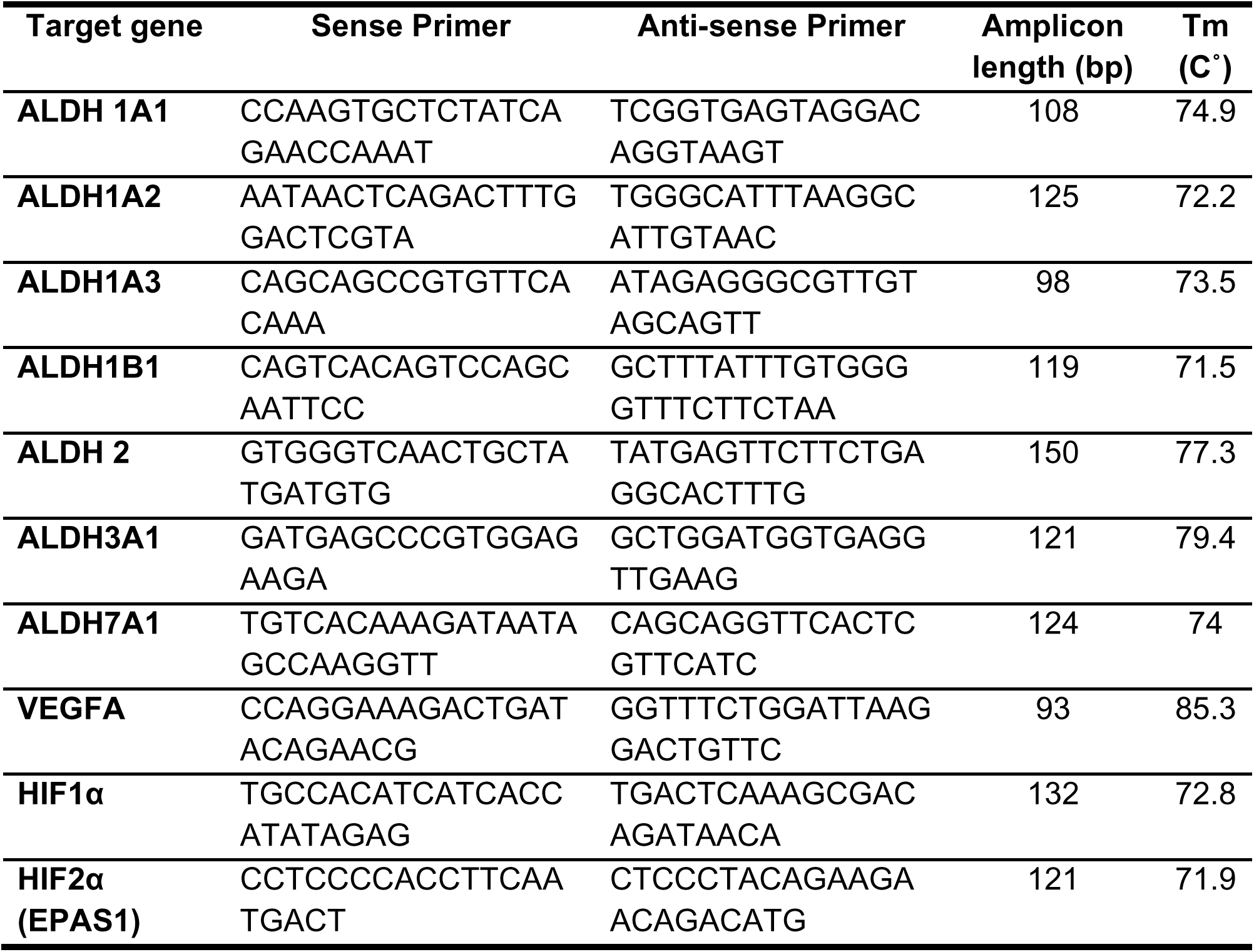
qRT-PCR primers. All primers were purchased from PrimerDesignTM.

### Western blotting analysis

Cell pellets were lysed in RIPA buffer, sonicated (3 × 10 s, power setting 10), and centrifuged at 13,200 × g for 15 min at 4°C. Protein samples were separated on 12% polyacrylamide gels and transferred to nitrocellulose membranes (GE Healthcare Life Sciences).

Membranes were incubated overnight at 4°C with primary antibodies diluted in 5% (w/v) non-fat milk in PBST: ALDH7A1 (1:20,000), LDH-A (1:5,000) (Abcam) and H2AX (1:1,000) (Cell Signalling Technologies). Protein detection was performed using HRP-conjugated secondary antibodies (Dako) and enhanced chemiluminescence (Roche). Membranes were then washed in PBST (3 × 5 min) and reprobed with an anti-actin antibody (1:80,000; Millipore), followed by a secondary antibody (1:10,000; Dako) as a loading control. Western blotting was also performed to assess the expression of ALDH1A1, ALDH1A2, ALDH1A3, and ALDH3A1 using antibodies listed in Appendix Table 2.

**Table 2:**
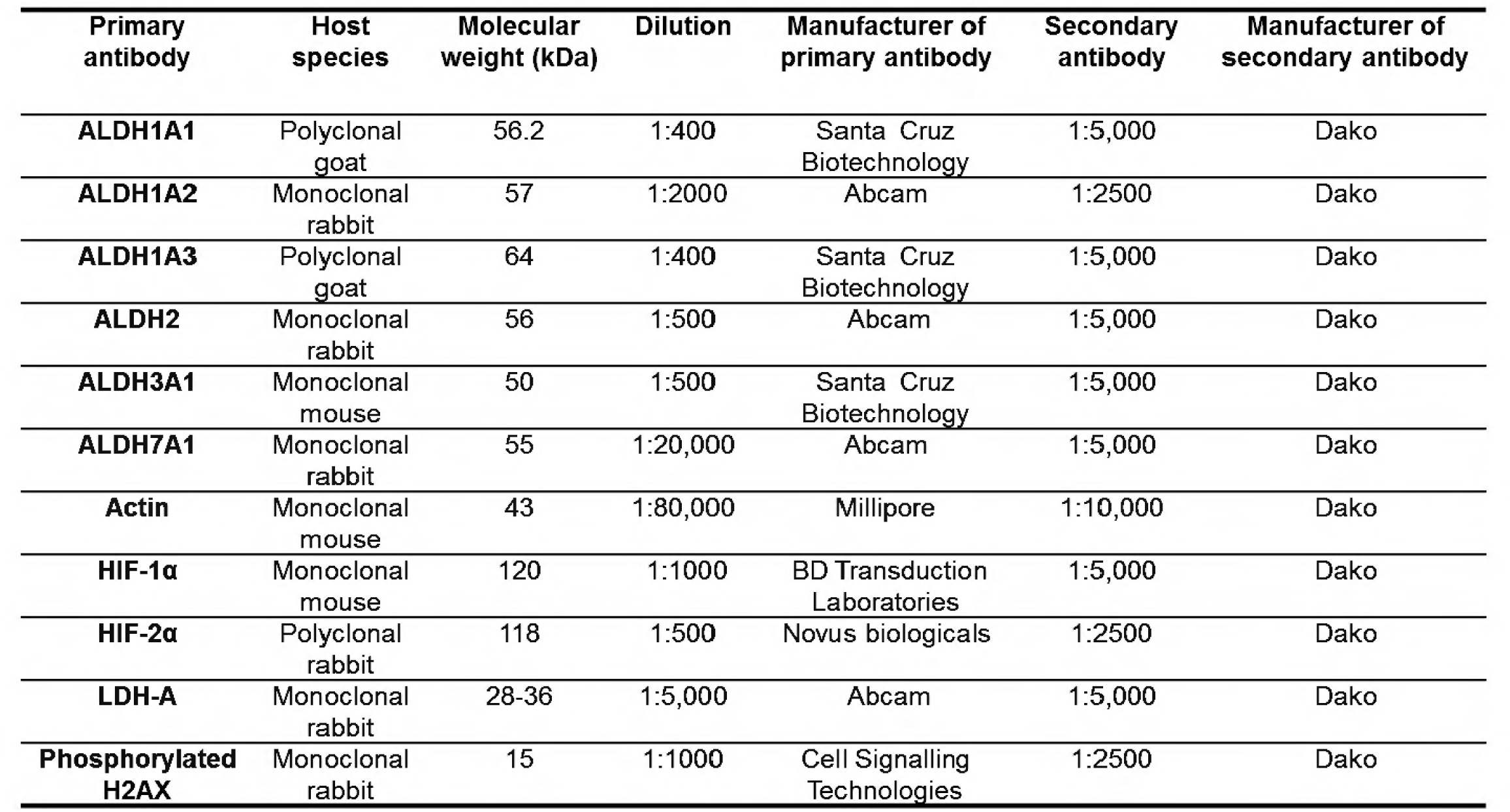
Primary and secondary antibodies for western blot.

### Spheroid formation assay

Spheroids were generated using the spinner flask culture method as previously described [27]. Briefly, cells were seeded in 250 mL spinner flasks at a density of 4 × 10⁶ cells in 100 mL complete RPMI medium and cultured at 37°C with continuous stirring at 60 rpm using a magnetic stirrer plate (Techne, Bibby Scientific Ltd., Stafford, UK). The culture medium was replenished every 2 days, and spheroid diameter was measured on alternate days from day 3 onwards. Spheroids reaching approximately 600 μm in diameter were harvested for RNA and protein extraction and subsequent analyses.

### Hypoxia detection

Hypoxic regions within spheroids were detected using the Hypoxyprobe™-1 Green Kit (HPI). Spheroids were incubated with pimonidazole (100 μM) for 2 h at 37°C, washed, embedded in OCT compound (Cryotek), snap-frozen using Cryo-Freeze Aerosol, and stored at −80°C. Frozen spheroids were cryosectioned at 5 μm and mounted on APES-coated slides. Sections were fixed in cold acetone for 10 min, washed, and blocked for 30 min in PBS containing 4% (v/v) FBS, 5% (w/v) non-fat milk, and 0.1% (v/v) Triton X-100. Pimonidazole adducts were detected by incubation with FITC-conjugated MAb1 antibody (1:150 dilution; supplied with the kit) for 2 h at 37°C. Nuclei were counterstained with DAPI (VECTASHIELD Mounting Medium with DAPI, Vector Laboratories). Fluorescence images were acquired using a Leica fluorescence microscope and Leica Application Suite Advanced Fluorescence (LAS AF) software.

### Selective trypsinisation of spheroids

Cells from distinct spheroid layers were isolated by sequential trypsinisation as previously described [28]. Briefly, spheroids were incubated with trypsin-EDTA for 2 min at room temperature with gentle agitation. Trypsinisation was stopped by adding 10 mL complete growth medium, and detached cells were separated from intact spheroids by sedimentation. The recovered cells were pelleted by centrifugation, snap-frozen on dry ice, and stored at −80°C until RNA or protein extraction. This process was repeated sequentially to remove successive cell layers until the spheroids were completely dissociated. Following each round of trypsinisation, the diameter of the remaining spheroid core was measured using a calibrated graticule mounted on a light microscope at 10× magnification.

For expression analyses, cells isolated from the first stripped layer were designated as the surface layer (SL). In contrast, cells obtained from the final viable layer adjacent to the necrotic core were designated as the hypoxic region (HR). In HT29 spheroids, the SL and HR corresponded to depths of 0-10.8 μm and 132-186 μm, respectively. In DLD-1 spheroids, the corresponding depths were 0-21 μm and 133-150.7 μm.

### Immunohistochemistry of spheroids

Spheroids were fixed in Bouin’s solution (Sigma) for 75 min at 37°C, stored in 70% ethanol, and processed for paraffin embedding [29]. Sections (5 μm) from the central region of the spheroids were deparaffinised, rehydrated, and stained with haematoxylin and eosin (H&E) for morphological assessment. For ALDH7A1 detection, sections underwent heat-induced antigen retrieval in citrate buffer, followed by blocking with normal goat serum (Vector Laboratories) for 30 min. Sections were incubated with anti-ALDH7A1 primary antibody (1:100) for 1 h, followed by a biotinylated goat anti-rabbit secondary antibody (1:200; Vector Laboratories) and avidin-biotin complex (Vectastain ABC kit). Immunoreactivity was visualised using DAB (Vector Laboratories), and sections were counterstained with Harris’ haematoxylin. Similar procedures were used to detect ALDH1A1, ALDH1A3, and ALDH3A1 (Appendix Table 3).

**Table 3:** Primary and secondary antibodies for IHC.

| **Primary antibody** | **Antigen retrieval** | **Blocking reagents** | **Dilution of primary antibody** | **Primary antibody incubation condition** | **Secondary antibody** |
| --- | --- | --- | --- | --- | --- |
| **ALDH1A1**<br>**(Polyclonal goat, Santa Cruz)** | 2 x 15 min<br>Citrate<br>Buffer | • 3% H<sub>2</sub>O<sub>2</sub> block<br>• Normal Rabbit Serum (1.5:100 in PBS for 30 min, Vector Laboratories) | 1:10 | 1h at 37°C | biotinylated rabbit anti goat antibody (1:200)<br>Vector Laboratories |
| **ALDH1A3**<br>**(Polyclonal goat, Santa Cruz)** | 2 x 15 min<br>Citrate<br>Buffer | • 3% H<sub>2</sub>O<sub>2</sub> block<br>• Normal Rabbit Serum (1.5:100 in PBS for 30 min, Vector Laboratories) | 1:10 | 1h at 37°C | biotinylated rabbit anti goat antibody (1:200)<br>Vector Laboratories |
| **ALDH3A1**<br>**(Monoclonal mouse, Santa Cruz)** | 2 x 15 min<br>Citrate<br>Buffer | • 3% H<sub>2</sub>O<sub>2</sub> block<br>• Normal Horse Serum (5:100 in PBS for 30 min, Vector Laboratories) | 1:10 | 1h at 37°C | biotinylated horse anti mouse antibody (1:200)<br>Vector Laboratories |
| **ALDH7A1**<br>**(Monoclonal rabbit, Abcam)** | 1 x 20 min<br>Citrate<br>Buffer | • 3% H<sub>2</sub>O<sub>2</sub> block<br>• Normal Goat Serum (1.5:100 in PBS for 30 min, Vector Laboratories) | 1:100 | 1h at room temperature | biotinylated goat anti rabbit antibody (1:200)<br>Vector Laboratories |

For generation of cell line-derived tumour xenografts, female Balb/C immunodeficient mice (aged six to eight weeks) were obtained from Envigo (Loughborough, UK) and injected subcutaneously with human CRC cell lines (HT29, HCT116, DLD-1, SW620 and COLO205). When the tumour size reached 500 mm^3^, mice were euthanised, and the tumours were excised, fixed in 10% formalin for 24 h, and processed for paraffin embedding before immunohistochemical (IHC) studies using the protocol for detecting ALDH expression. This investigation was conducted in accordance with ethical standards approved by the Animal Welfare Ethics Review Board at the University of Bradford and with the UK National Cancer Research Institute Guidelines for the Welfare of Animals [30]. Throughout the study, all mice were housed in air-conditioned rooms in facilities approved by the United Kingdom Home Office to meet all current regulations and standards, and received the Teklad 2018 diet (Envigo, Blackthorn, UK) and water *ad libitum*. All procedures were carried out under a Project Licence (PPL 40/3670) issued by the UK Home Office in accordance with government legislation.

### Induction of HIF using cobalt chloride (CoCl_2_)

Cobalt chloride (CoCl₂), a chemical inducer of HIF-1α under normoxic conditions [31], was used to investigate whether ALDH7A1 expression was regulated by HIF-1α. To determine non-toxic concentrations, DLD-1 and HT29 cells (2 × 10³ cells/well) were seeded in 96-well plates and treated with CoCl₂ (10-500 μM) for 24 h. The medium was then replaced with fresh medium, and cells were incubated for a further 72 h. Cell viability was assessed using the MTT assay as previously described [32]. Briefly, cells were incubated with MTT solution (0.5 mg/mL) for 4 h, after which the medium was removed, and formazan crystals were dissolved in DMSO. Absorbance was measured at 540 nm using a microplate reader (Thermo Electron Corporation).

For analysis of HIF-1α and ALDH7A1 expression, DLD-1 and HT29 cells (2 × 10⁵ cells) were seeded into T75 flasks and treated with CoCl₂ for 24 h. DLD-1 cells were exposed to 100 or 150 μM CoCl₂, while HT29 cells were treated with 200 or 300 μM CoCl₂. Cells were then harvested for protein extraction, and HIF-1α and ALDH7A1 expression was analysed by Western blotting as described above.

### siRNA transfections

siRNA duplexes targeting HIF1α, HIF2α, ALDH1A3, ALDH3A1, and ALDH7A1 were designed, synthesised, and validated by Ambion/Life Technologies (Appendix Table 4). DLD-1 cells were seeded at a density of 2.75 × 10⁵ cells per T25 flask and incubated for 24 h before transfection. Transfections were performed as previously described [33]. Briefly, siRNA was diluted in Opti-MEM (Gibco) and complexed with a transfection reagent before being added to cells at a final siRNA concentration of 20 nM. Liposome-only and mock-transfected cells served as controls. Following 4 h incubation, an equal volume of 2× RPMI medium was added. For hypoxic experiments, pre-equilibrated hypoxic medium was used, and cells were maintained under 0.1% O₂. QRT-PCR and Western blotting were used to assess knockdown efficiency. Gene expression changes in siRNA-transfected cells were compared with mock-transfected and liposome control cells.

**Table 4:**
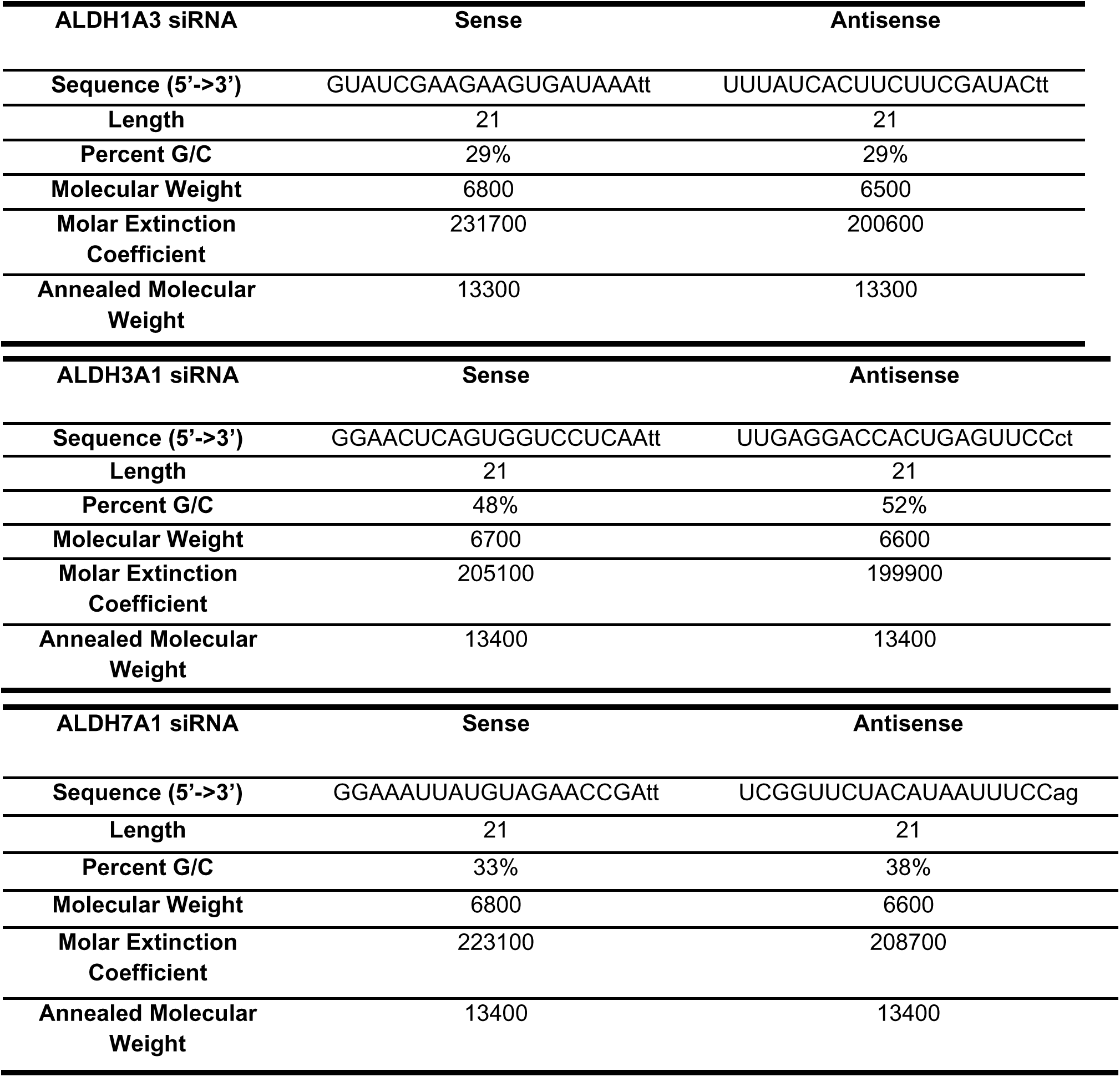
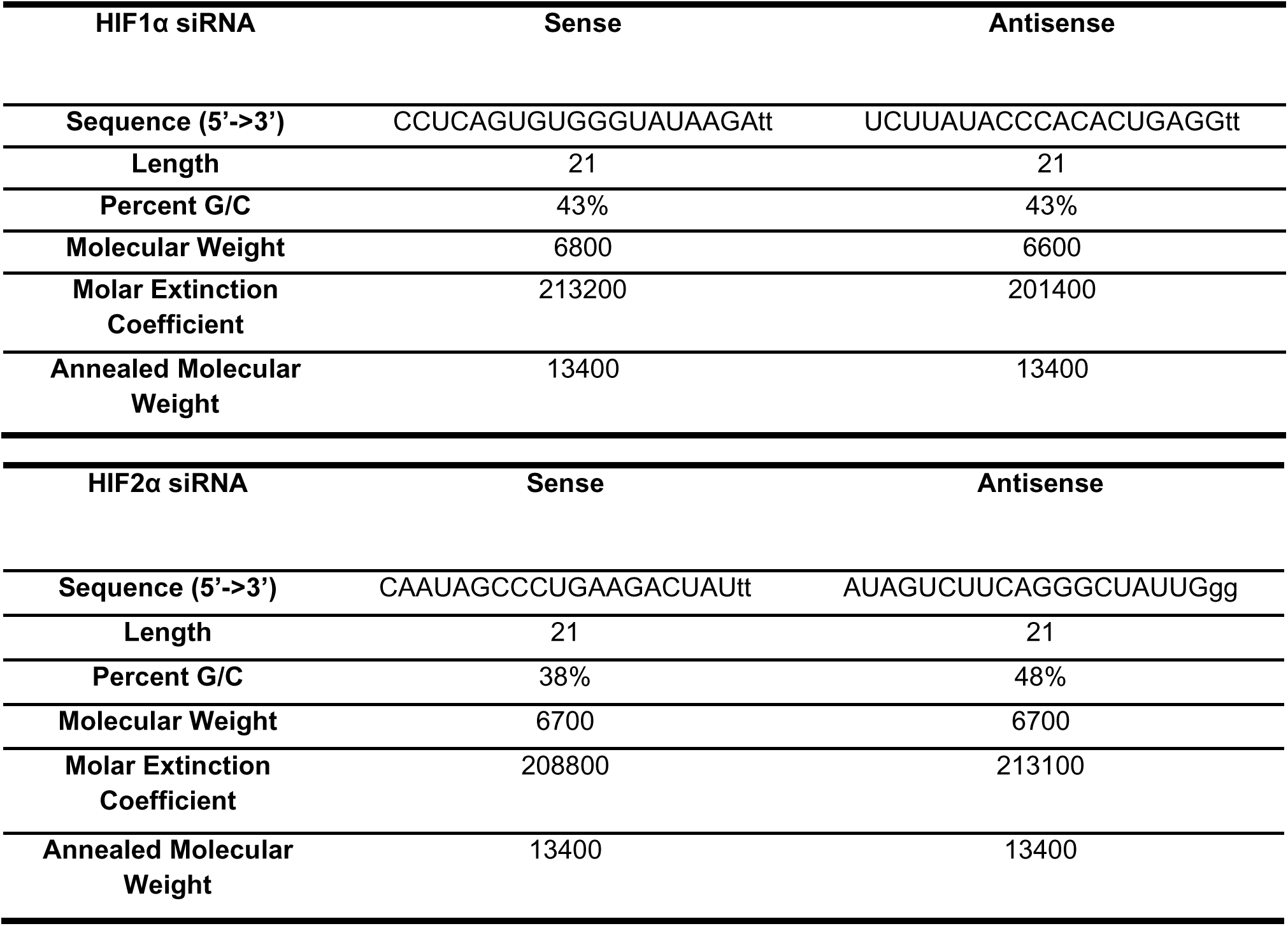
siRNAs for target genes were purchased from Ambion/Life Technologies.

### Detection of reactive oxygen species (ROS)

Intracellular ROS levels were measured 72 h post-transfection using 6-carboxy-2′,7′-dichlorodihydrofluorescein diacetate (carboxy-H2DCFDA; Fisher Scientific). Cells were harvested and incubated with 5 μM carboxy-H2DCFDA in phenol red-free RPMI for 30 min at 37°C, with gentle agitation to prevent cell attachment. Cells were then washed and resuspended in PBS and analysed by flow cytometry using the FL1 channel. Cells treated with 250 μM H_₂_O_₂_ following 48 h of transfection served as a positive control for ROS generation.

### Cell proliferation and viability

To evaluate the role of ALDH isoforms in cell proliferation and viability, DLD-1 cells were seeded at a density of 2.75 × 10⁵ cells per T25 flask and transfected with siRNAs as described above. At 24, 48, and 72 h post-transfection, cells were harvested and stained with trypan blue. Live and dead cells were counted using a haemocytometer, and cell proliferation and viability were determined from the total number of viable and non-viable cells.

### Drug cytotoxicity assay

DLD-1 cells were seeded in 6-well plates at a density of 1.1 × 10⁵ cells per well and transfected with siRNA as described above. After 48 h, cells were treated with oxaliplatin (75 μM), irinotecan (75 μM), or 5-fluorouracil (5-FU; 100 μM) for a further 48 h. Vehicle control cells received 0.1% DMSO. Cell viability was assessed by trypan blue exclusion, and the number of viable cells was determined using a haemocytometer.

To assess the effect of hypoxia, the experiment was repeated with ALDH7A1 siRNA-transfected cells under hypoxic conditions. Cell survival was calculated as:

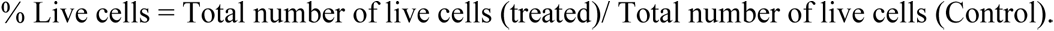

### Statistical analysis

Data are presented as the mean ± standard deviation (SD) from at least three independent experiments. Statistical analyses were performed using a two-tailed Student’s *t*-test to compare group means. Differences were considered statistically significant at *p* < 0.05.

## Results

### Hypoxia induces the expression of selective ALDH isoforms in CRC cells

It has been demonstrated in various cancers that ALDH isoforms, such as ALDH1A1 and ALDH1A3, are upregulated in hypoxic niches and correlate with stemness [34,35]. However, in CRC, no comprehensive studies have profiled the expression signatures of ALDH isoforms in response to hypoxic conditions, specifically recapitulating the acute or chronic hypoxia observed in solid tumours. To address this gap, transcript and protein level expression of a set of seven selected ALDH isoforms (1A1, 1A2, 1A3, 1B1, 2, 3A1 and 7A1) that have previously been linked to cancer pathogenesis were evaluated in four established CRC cell lines (DLD-1, HCT116, HT29 and SW480) under hypoxic (0.1%) and normoxic conditions. qRT-PCR analysis of the selected target genes in CRC cell lines under normoxic conditions revealed a similar trend in the basal level expression patterns of ALDH isoforms, with increased expression of ALDH1A3, 1B1, 2, and 7A1 compared to ALDH1A1 and 1A2 (Figure 1A-D, left panel). In contrast to other CRC cell lines examined, HT29 cells showed lower differential expression of the isoforms, with ALDH1A2 exhibiting the lowest. Exposure of the cells to hypoxia demonstrated a significant impact on ALDH expression, which appeared cell line-specific, although ALDH1A2 was observed to be consistently induced in all the cell lines investigated (Figure 1A-D, right panel). The expression of ALDH1A1 was significantly upregulated in HT29 and SW480 cells under hypoxic conditions, while DLD1 and SW480 cells showed an increased expression of ALDH1A3. Exposure to hypoxia also upregulated ALDH3A1 and ALDH1B1 expression in DLD-1 and HT-29 cells, respectively.

**Figure 1.**
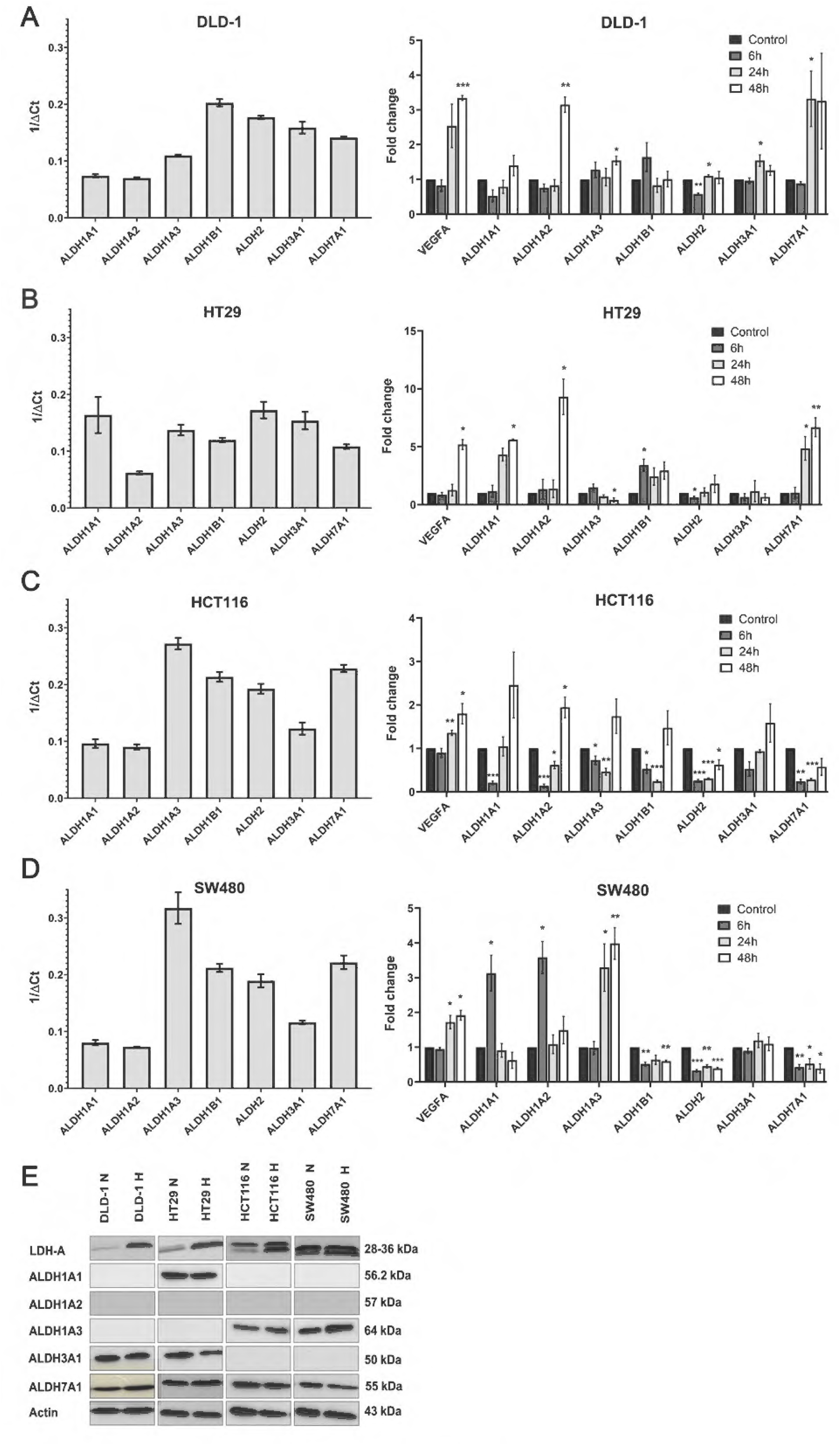
Expression profiling and hypoxia-regulated expression of ALDH isoforms in CRC cell lines. Expression of ALDH1A1, ALDH1A2, ALDH1A3, ALDH1B1, ALDH2, ALDH3A1 and ALDH7A1 mRNAs was assessed by qRT-PCR in DLD-1 (A), HT29 (B), HCT116 (C) and SW480 (D) monolayer cells under normoxic (control) and hypoxic conditions (0.1% O₂ for 6 h, 24 h and 48 h). Left panels show mRNA abundance expressed as 1/ΔCt, where ΔCt = Ct (target gene) − Ct (β-actin). Right panels show fold change relative to normoxic control, with VEGFA included as a positive control. Values represent the mean of three independent experiments, and error bars indicate SD. P values: * p<0.05, ** p<0.01, *** p<0.001. (E) Western blot analysis of ALDH in CRC cell lines under normoxic and hypoxic conditions (48 h exposure to 0.1% O2). LDH-A was used as a positive control for hypoxia induction, and β-actin was used as a loading control.

Interestingly, ALDH7A1 was significantly upregulated in HT29 cells (approximately 5-and 7-fold after 24 h and 48 h of hypoxia exposure, respectively) and in DLD-1 cells (3-fold after 24 h of hypoxia exposure).

Further, the protein expression levels of the ALDH isoforms were analysed under similar conditions. ALDH1B1 and 2 were not considered for the protein analysis because these isoforms were not significantly affected by exposure to low-oxygen tension over 48 h. Western blotting analysis of the isoforms revealed that ALDH1A1 protein was expressed only in HT29 cells, whereas ALDH1A2 was not detectable in any of the cell lines examined (Figure 1E). ALDH1A3 was expressed in HCT116 and SW480 cells, while ALDH3A1 was present only in DLD-1 and HT29. In contrast to all these isoforms, ALDH7A1 was expressed at the protein level in all four CRC cell lines. Following 48 h of exposure to hypoxia (0.1% O_2_), ALDH7A1 was found to be upregulated in DLD-1 and HT29 cells, while an increased expression of ALDH1A3 was observed in HCT116 and SW480 cells, consistent with the mRNA expression pattern. Altogether, these findings indicated a cell line-specific impact of hypoxia on the gene-and protein-level expression of ALDH isoforms.

### ALDH7A1 is upregulated in the hypoxic region of multicellular spheroids (MCS)

Next, the expression of these isoforms in MCS as a three-dimensional model system was assessed, as it more closely resembles the functional and microenvironmental properties of tumours growing *in vivo* [36]. For this purpose, DLD-1 and HT29 cells were grown in spinner flasks to generate spheroids, which were observed following three days of culture (Figure S1). The cellular heterogeneity within an MCS replicated the histomorphological properties of the tumour, wherein an external layer displaying a gradient of proliferating cells surrounds the centrally located quiescent cells, which enclose the hypoxic core. Cells in the hypoxic core of a spheroid are comparable to the less proliferative or dormant cells present in an avascular tumour core *in vivo,* which display stem cell properties and are refractory to the current anti-cancer therapies. To evaluate ALDH isoform expression across distinct spheroid regions, cells from the surface layer (SL) and hypoxic region (HR) of HT29 and DLD-1 multicellular spheroids were isolated by sequential trypsinisation and analysed for gene and protein expression. The SL consisted of cells with access to oxygen and nutrients, whereas the HR contained oxygen-and nutrient-deprived cells.

Gene expression analysis showed that ALDH7A1 was the only isoform consistently upregulated in both cell lines within the spheroids. In HT29 and DLD-1 spheroids, ALDH7A1 expression increased in both the SL and HR compared with normoxic monolayer cultures (Figure 2). In the hypoxic region (HR), ALDH7A1 expression was increased by 8.5-fold in HT29 spheroids and 3.0-fold in DLD-1 spheroids relative to monolayer controls. Although the increase in ALDH7A1 expression in HR compared to SL did not reach statistical significance, likely due to biological variability, the consistent upregulation observed in both cell lines prompted further investigation of this isoform. HT29 spheroids also exhibited significant upregulation of ALDH1A1, ALDH1A2, and ALDH1B1 mRNA expression. However, no marked differences in the expression of these isoforms were observed between the surface layer (SL) and HR, except for ALDH1A1, which was enriched in the HR of DLD-1 spheroids (Figure 2A). In addition, ALDH2 expression was elevated in HR cells from both cell lines.

**Figure 2.**
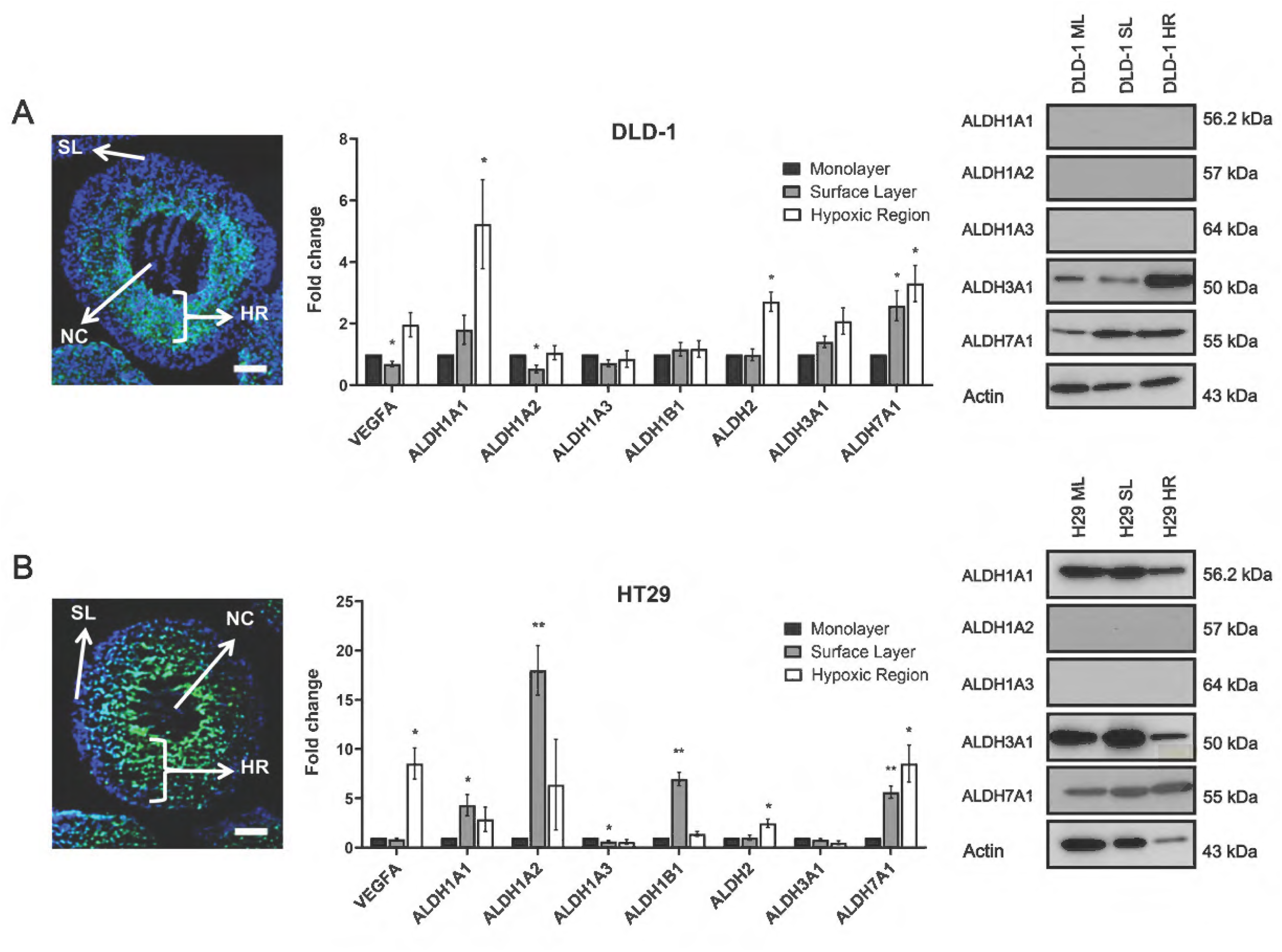
ALDH mRNA and protein expression in DLD-1 (A) and HT2G (B) multicellular spheroids (MCS). Necrotic core (NC), surface layer (SL) and hypoxic regions (HR) were identified by pimonidazole (green) and DAPI staining (blue). Scale bar = 100 μm at 10x objective lens. ALDH isoform mRNA expression was analysed using qRT-PCR in the monolayer, surface layer (SL) and hypoxic regions (HR) of the MCS, with VEGFA as a positive control and β-actin as an internal control. Protein expression was assessed using western blot, with actin as the internal control protein. Values are the mean of three independent experiments, and error bars represent SD. P values: * p<0.05, ** p<0.01.

Western blot analysis was performed to determine whether changes in mRNA expression were reflected at the protein level. In HT29 spheroids, ALDH7A1 protein expression increased by 1.9-fold in the surface layer (SL) and 4.6-fold in the HR relative to monolayer cultures (Figure 2B, right panel). Similarly, ALDH7A1 expression was elevated in DLD-1 spheroids, showing a 2.7-fold increase in the SL and a 3.0-fold increase in the HR. In addition, ALDH1A1 and ALDH3A1 protein expression were increased in the HR of HT29 (1.8-fold) and DLD-1 (1.84-fold) spheroids, respectively. The variable protein expression of the isoforms across different layers of the MCS warranted further exploration of the distribution and cellular localisation of these selected ALDH isoforms. Immunohistochemistry was therefore performed to assess the spatial expression patterns of selected ALDH isoforms (ALDH1A1, ALDH1A3, ALDH3A1, and ALDH7A1) in MCS; n =2: HT29 and DLD-1), cell line-derived xenograft (CDX) models (n =5: HT29, DLD-1, HCT116, SW620, and COLO205), and primary clinical colorectal cancer (CRC) specimens (n =50).

ALDH1A1 exhibited predominantly cytoplasmic staining throughout the spheroids, with stronger expression in the deeper hypoxic regions than in the surface layers. Positive ALDH1A1 staining was also detected in some cells within the necrotic core (Figure 3A). In contrast, ALDH1A3 expression was not detected in either HT29 or DLD-1 spheroids, consistent with the Western blot results. ALDH3A1 displayed both cytoplasmic and nuclear staining in HT29 and DLD-1 spheroids, with positive nuclear staining evident within the necrotic core (Figure 3A). Increased ALDH3A1 expression was observed in the hypoxic region of DLD-1 spheroids but not in HT29 spheroids, consistent with the western blot findings.

**Figure 3.**
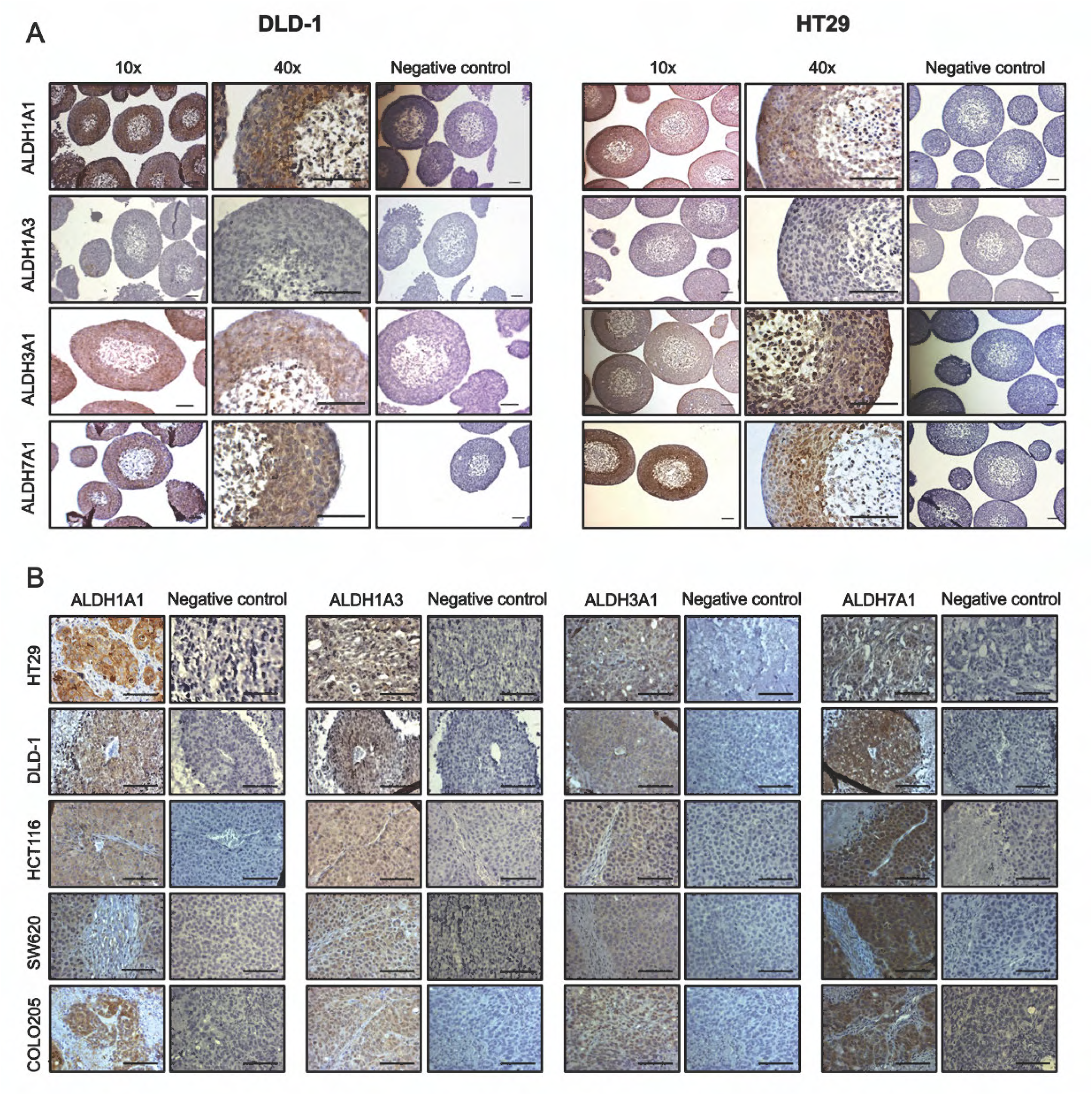
ALDH isoform expression in CRC multicellular spheroids and colon cancer xenografts. (A) Immunohistochemical staining for ALDH1A1, ALDH1A3, ALDH3A1 and ALDH7A1 in DLD-1 (left panel) and HT29 (right panel) multicellular spheroids (MCS), shown at x10 and x40 magnification with negative controls for each isoform. (B) Immunohistochemical staining for ALDH1A1, ALDH1A3, ALDH3A1 and ALDH7A1 in colon cancer xenografts derived from HT29, DLD-1, HCT116, SW620 and COLO205 cells. Brown colour indicates positive staining and ALDH expression. Scale bar = 100 µm.

In contrast, elevated cytoplasmic and nuclear ALDH7A1 expression was detected in the hypoxic region of MCS from both DLD-1 and HT29 cell lines, which gradually decreased with increasing distance from the peripheral surface layers. Some cells in the necrotic core of both MCS models were also stained positively for ALDH7A1. Cell line-derived xenografts exhibited similar cytoplasmic and nuclear staining patterns for all ALDH isoforms analysed by immunohistochemistry (Figure 3B). Given the pronounced upregulation of ALDH7A1 in hypoxic regions and its established role in protection against oxidative stress [25], this isoform was selected for further investigation to determine whether its increased expression represents an adaptive cytoprotective response to hypoxic stress.

### ALDH7A1 co-expresses with GLUT-1 in the hypoxic fractions of human CRC primary tumours

Given that ALDH7A1 expression was consistently elevated under hypoxic conditions in CRC cell lines and multicellular spheroids, its association with hypoxia was further investigated in primary CRC specimens by assessing its co-expression with the hypoxia marker GLUT1. Analysis of a tissue microarray containing archived CRC patient samples revealed significantly higher expression of both ALDH7A1 and GLUT1 in tumour tissues compared with matched adjacent non-tumour tissues. Intensity of GLUT1 expression was elevated across all CRC stages examined (I, IIa, IIb, III, and IIIb) relative to adjacent normal tissue (Figure 4A). In contrast, ALDH7A1 expression was significantly increased only in stage I tumours (*p* < 0.05). Representative immunohistochemical images demonstrating ALDH7A1 expression in both normoxic and hypoxic regions of CRC tissue are shown in Figure 4.

**Figure 4.**
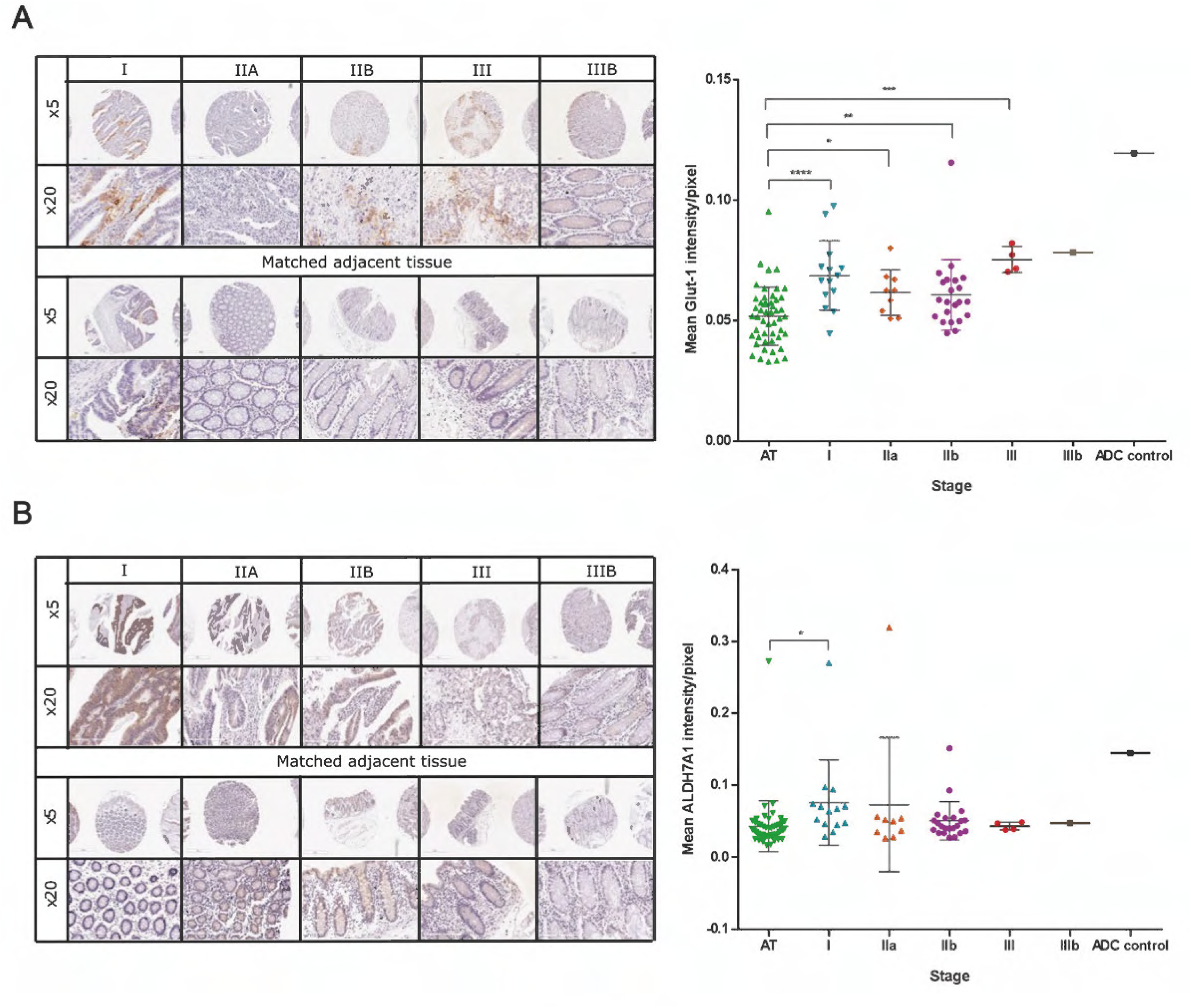
ALDH7A1 and GLUT1 expression in colon adenocarcinoma tissue microarray (TMA). Left panels show representative immunohistochemical images of GLUT1 (A) and ALDH7A1 (B) expression across tumour stages I, IIA, IIB, III and IIIB and matched adjacent tissue, shown at 5× and 20× magnification. Images were acquired using Aperio Leica software. Brown staining indicates positive expression. Right panels show mean expression of GLUT1-positive (A) and ALDH7A1-positive (B) across tumour stages and adjacent tissue (AT) measured as intensity/pixel. Each data point for cancer tissue is an average taken from 2 different cores of each patient case. 50 cases were analysed: 2 cancer cores and 1 adjacent tissue, totalling 150 individual images. Data analysis was performed using QuPath software, and statistical test applied using GraphPad Prism. Statistical differences were compared between each cancer stage and the mean of adjacent tissue using a t-test. P values: * p<0.05, ** p<0.01, *** p<0.001.

### Upregulation of ALDH7A1 by hypoxia occurs through a HIF-1-independent mechanism

It is well established that tumour cells adapt to oxygen deprivation through the stabilisation of hypoxia-inducible factors (HIFs) [37] with HIF-1α as a key regulator of the adaptive cellular response of tumour cells to hypoxia [38]. To assess whether HIF-1α mediated the elevated expression of ALDH7A1 under hypoxic conditions, CRC cells were treated with cobalt chloride (CoCl₂), a hypoxia-mimetic agent that stabilises HIF-1α under normoxic conditions [39]. DLD-1 and HT29 cells exhibited differential sensitivity to CoCl₂ treatment in the viability assay (Figure S2). Based on the dose-response profiles, concentrations of 100 and 150 μM were selected for DLD-1 cells, and 200 and 300 μM for HT29 cells, to achieve HIF-1α stabilisation while maintaining acceptable cell viability. CoCl₂ treatment resulted in robust HIF-1α expression in both cell lines; however, no corresponding increase in ALDH7A1 expression was observed (Figures 5A and 5B). These findings suggest that HIF-1α does not directly mediate ALDH7A1 upregulation under hypoxic conditions.

**Figure 5.**
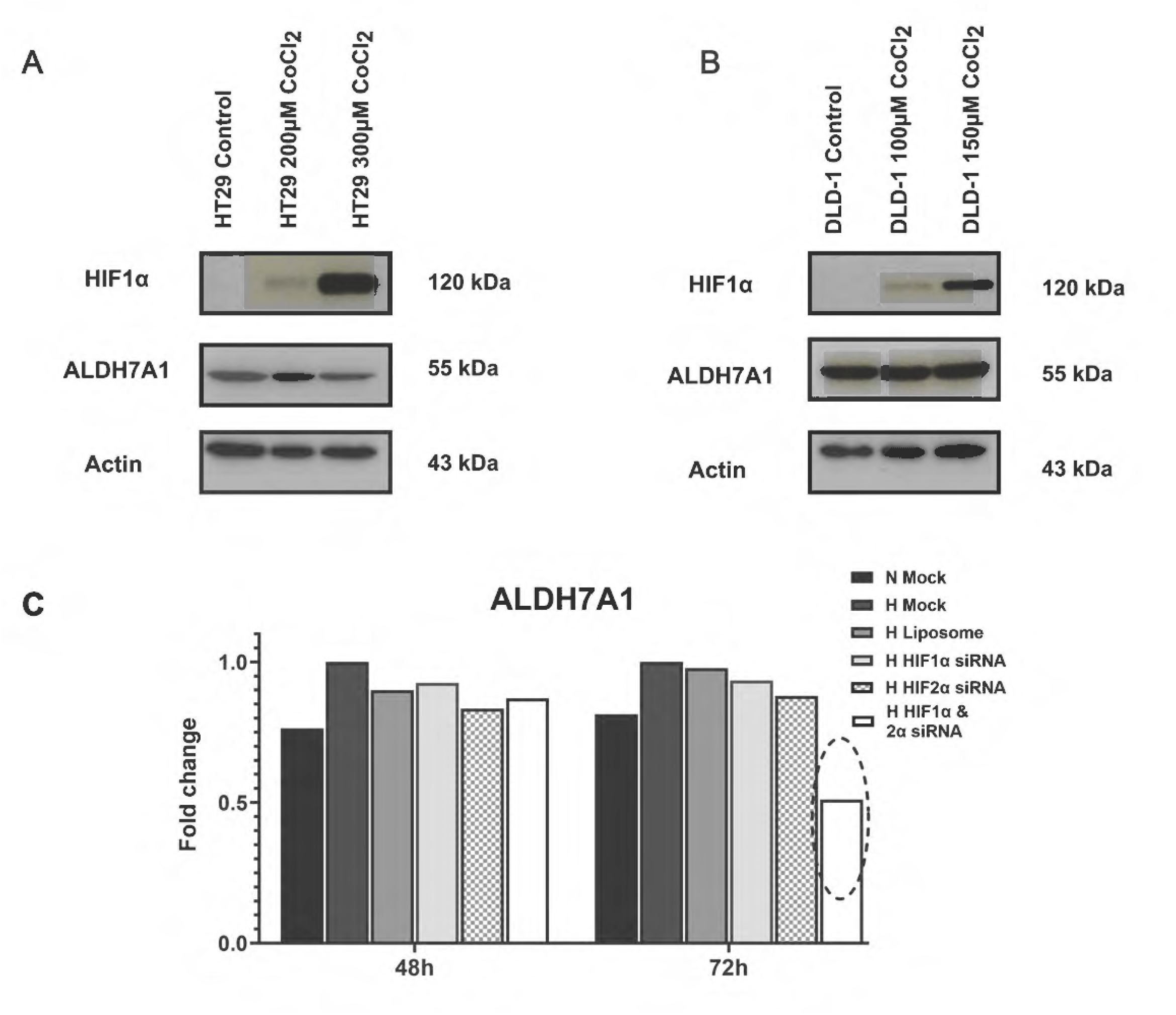
Regulation of ALDH7A1 by HIF. Western blot analysis of HIF1α and ALDH7A1 protein expression upon treatment with CoCl_2_ (concentration range from 100 µM to 300 µM) in HT29 cells (A) and DLD-1 cells (B) under normoxic conditions. (C) Expression of ALDH7A1 after HIF knockdown. ALDH7A1 mRNA expression measured using qRT-PCR after 48 h and 72 h of HIF1α, HIF2α or dual HIF1/2α siRNA-mediated knockdown, under hypoxic conditions. Normoxic and hypoxic mock served as basal expression controls under normoxia and hypoxia, respectively.

To further confirm the absence of any regulatory role of HIF subunits in hypoxia-induced upregulation of ALDH7A1, single and dual knockdown studies using small interfering RNAs (siRNAs) against HIF-1α and/or HIF-2α were carried out under hypoxic conditions, and their effect on ALDH7A1 expression was evaluated. Significant and specific reduction in HIF-1α and HIF-2α gene expression was achieved after 48 h and 72 h of single or dual siRNA transfection (Figure S3). However, consistent with the previous results, the mRNA analysis of ALDH7A1 in HIF-knockdown cells revealed that isoform expression was unaffected even after 48 h or 72 h of single HIF knockdown. Nevertheless, a dual knockdown of HIF-1α/2α resulted in a 50% reduction in ALDH7A1 mRNA expression after 72 h (Figure 5C), with a reduction of less than 22% at the protein level. These results suggested that the upregulation of ALDH7A1 expression under hypoxic conditions is independent of HIF1α/2α regulation.

### Probing the functional roles of ALDH7A1 in colorectal cancer using siRNA knockdown

To further elucidate the functional role of ALDH7A1 in cancer progression and drug resistance, ALDH7A1 knockdown studies were performed in the DLD-1 cell line under normoxic and hypoxic conditions. The DLD-1 cell line was chosen for this study because it exhibited the highest ALDH7A1 expression under hypoxia and showed upregulated ALDH7A1 protein in the hypoxic region of MCS and xenografts. Cell images taken at 24 h, 48 h, and 72 h following siRNA transfection showed no obvious phenotypic differences among siRNA-transfected cells, liposome controls, and mock cells under normoxic and hypoxic conditions (data not shown).

To evaluate the specificity of target siRNA and to assess the possibility of crosstalk between different ALDH isoforms, analysis of the effect of ALDH isoform knockdown on other selected members (ALDH1A3 and ALDH3A1) of the ALDH family was also carried out. Knockdown of ALDH3A1 and ALDH7A1 isozymes was successfully achieved, resulting in the abolishment of up to 70% of target ALDH expression in DLD-1 cells at both the mRNA and protein levels under normoxic conditions (Figure S4). The efficiency of ALDH1A3 knockdown was evaluated only at the gene level, as ALDH1A3 protein was not detectable in DLD-1 cells.

Interestingly, silencing of ALDH7A1 resulted in consistent upregulation of ALDH3A1 mRNA and protein expression (Figure 6A-C), suggesting a potential regulatory interplay among ALDH isoforms and that 3A1, directly or indirectly, can influence or compensate for the expression of 7A1. Knockdown studies were also conducted under hypoxic conditions, and the results showed that siRNAs suppressed expression at both the mRNA and protein levels (Figure S5). As in normoxia, ALDH7A1 knockdown under hypoxic conditions also resulted in upregulation of ALDH3A1, particularly after 48 h and 72 h of siRNA transfection, supporting crosstalk between these two isoforms (Figure 6D-F). Accordingly, dual knockdown of ALDH3A1 and 7A1 was performed under normoxic and hypoxic conditions. A reduction in the mRNA and protein levels of these isoforms was observed under both conditions (Figure S6). ALDH7A1/3A1 single-or dual-knockdown cells were further evaluated to elucidate the biological importance of these isoforms in mediating CRC progression.

**Figure 6.**
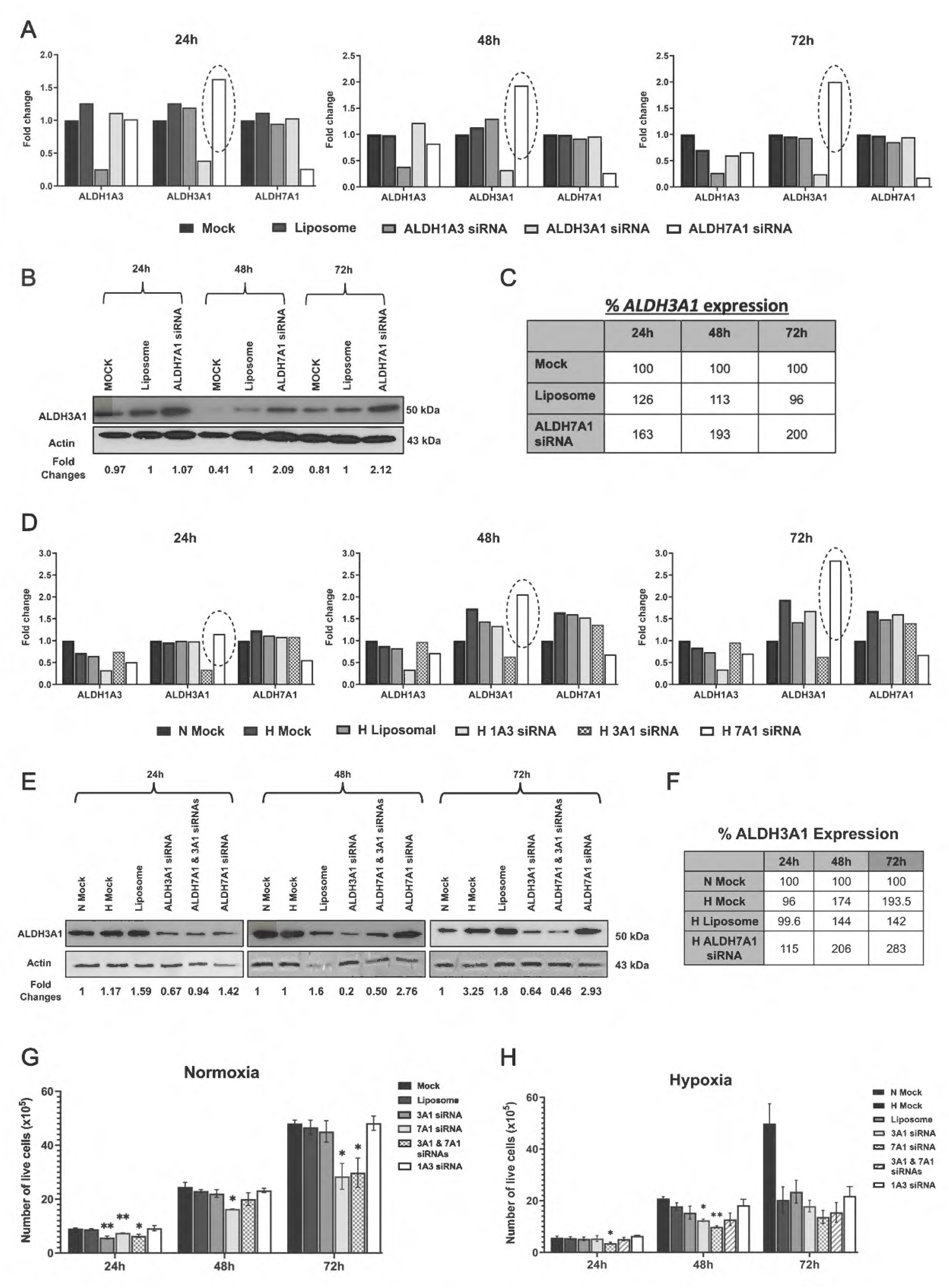
Characterising functional roles of ALDH isoforms through siRNA knockdown under normoxia and hypoxia. DLD-1 cells were transfected with ALDH (1A3, 3A1, and 7A1) siRNAs for 24 h, 48, and 72 h. (A, D) qRT-PCR analysis of ALDH1A3, 3A1 and 7A1 mRNA expression under normoxic (A) and hypoxic (D) conditions. Values expressed as fold change relative to normoxic mock control. Dashed circles highlight consistent upregulation of ALDH3A1 following ALDH7A1 siRNA transfection. (B, E) Western blot analysis of ALDH3A1 protein expression in ALDH7A1 siRNA-transfected cells under normoxia and hypoxia, respectively. Actin was used as a loading control, with relative fold changes relative to liposome control indicated below. (C, F) Quantification of ALDH3A1 gene expression in ALDH7A1 siRNA-transfected cells relative to mock control, under normoxic and hypoxic conditions, respectively. (G, H) Total live cell number in DLD-1 cells transfected with ALDH isoform siRNAs under normoxia and hypoxia at 24 h, 48 h and 72 h, assessed by trypan blue assay. Live cell number compared to N mock in normoxia (G) and H mock in hypoxia (H). Values represent the mean of three independent experiments, and error bars indicate SD. P values: * p<0.05, ** p<0.01.

### ALDH7A1 regulates cell proliferation in CRC under normoxic and hypoxic conditions

ALDH7A1 is involved in mediating cell growth and enhancing clonogenicity in prostate cancer [40]. To test whether ALDH7A1 may have a similar function in CRC, cell proliferation and survival were evaluated in ALDH7A1-knockdown cells using the trypan blue exclusion assay. Cells transfected with siRNA against ALDH7A1 proliferated at a significantly slower rate with fewer live cells at all time points under normoxia (*p* = 0.007, 0.02 and 0.02 for 24 h, 48 h and 72 h) (Figure 6G). Similar results were observed with the ALDH3A1 and ALDH7A1 dual knockdown cells, where the reduction in cell number was presumed to be due to reduced ALDH7A1 expression, as ALDH3A1 siRNA-transfected cells showed less cell number only at 24 h of transfection relative to mock cells (*p* = 0.007). No significant differences in the number of dead cells were detected between groups, indicating that the effects of ALDH7A1 depletion are more likely attributable to reduced cell proliferation than increased cell death.

Under hypoxic conditions, the rate of cell proliferation was much lower than in normoxia, consistent with many other studies showing the detrimental effects of hypoxia on cell proliferation [41]. ALDH3A1 siRNA-transfected cells showed fewer live cell numbers only after 48 h of transfection (*p* = 0.021). ALDH7A1 siRNA-transfected cells also had a reduced total number of live cells at 24 h and 48 h post-transfection (*p* = 0.017 and 0.010, respectively) compared to hypoxic mock control cells, supporting a regulatory role for this isoform in DLD-1 cell proliferation (Figure 6H).

### ALDH7A1 knockdown augments the generation of ROS

Although hypoxia induced ALDH7A1 expression, the mechanism by which this isoform contributes to cellular adaptation under hypoxic stress remained unclear. Emerging evidence suggests that members of the ALDH family, including ALDH3A1 and ALDH7A1, possess antioxidant functions. In particular, ALDH3A1 has been reported to protect cells against reactive oxygen species (ROS)-induced oxidative damage, suggesting that ALDH7A1 may exert a similar cytoprotective role under hypoxic conditions [25,42]. The detoxification capacity of ALDHs has also been suggested to be an important factor governing CSC longevity and to protect them against oxidative insults that are markedly increased in cancer [43,44]. However, no reports describe the role of ALDH7A1 in cancer under normal basal growth conditions, in the absence of external oxidative stress.

The generation of ROS in ALDH1A3/3A1/7A1 knockdown cells was detected by flow cytometry. Hydrogen peroxide (H_2_O_2_) was used as a positive control for ROS induction. Knockdown of ALDH1A3 reduced ROS in DLD-1 cells relative to liposome controls. At the same time, inhibition of ALDH3A1 had no effect, contrary to its previously reported function as a ROS scavenger [45] (Figure 7A and 7B). Knockdown of ALDH7A1 increased ROS under normoxic conditions compared to control cells, supporting the antioxidant properties of ALDH7A1 in DLD-1 cells. A similar finding was observed in the co-transfected cells, although no apparent effect was observed with ALDH3A1 knockdown alone. As elevated ROS levels can induce DNA damage, including double-strand DNA breaks that may ultimately lead to cell death [46], the effect of ROS on phosphorylated histone H2AX (γH2AX), a well-established marker of DNA double-strand breaks, was investigated [47]. However, no major difference in phosphorylated H2AX expression was observed for ALDH7A1 knockdown cells (Figure 7C).

**Figure 7.**
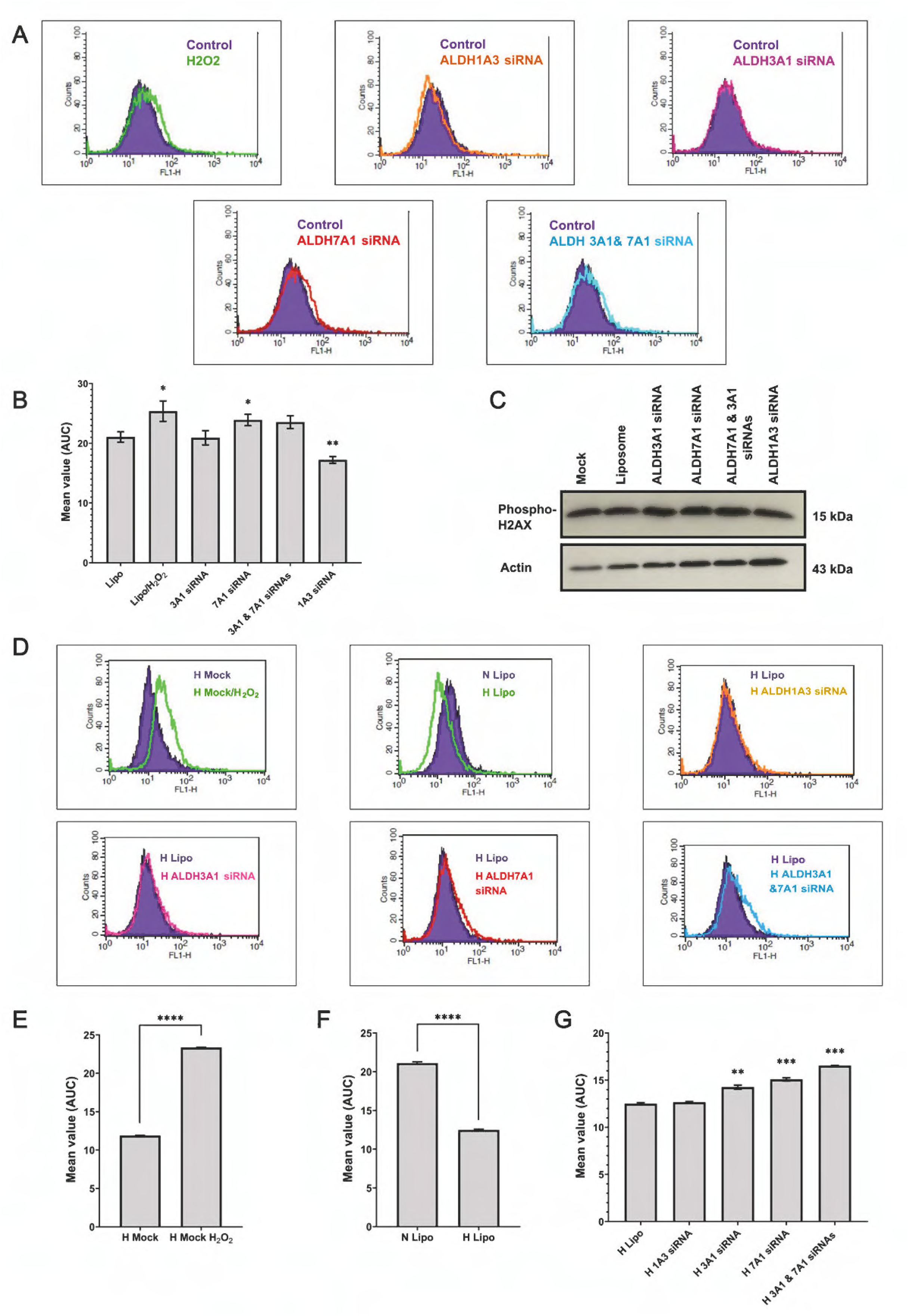
Detection of reactive oxygen species (ROS) generation in DLD-1 siRNA-transfected cells after 72 h under normoxic (A-C) and hypoxic conditions (D-G). (A, D) ROS generation curves under normoxic and hypoxic conditions, respectively. Geometric mean values of area under the curve (AUC) from ROS detection in DLD-1 cells after knockdown under normoxic (B) and hypoxic conditions (E-G). Values represent the mean of three independent experiments and error bars indicate SD. P values: * p<0.05, ** p<0.01, *** p<0.001, ****p<0.0001. (C) Western blot analysis of phosphorylated H2AX protein expression in siRNA knockdown samples in normoxic conditions.

Generation of ROS in a hypoxic TME remains a subject of debate [11,48]. As oxygen availability is a key determinant of cellular redox status, alterations in oxygen tension can markedly influence ROS production. In the present study, hypoxic cells exhibited lower ROS levels than their normoxic counterparts, consistent with previous reports [48,49]. Comparison between hypoxic knockdown samples and liposome control showed no major difference in ROS generation between ALDH1A3 knockdown (Figure 7D-F). ALDH3A1 knockdown significantly increased ROS generation (*p*=0.002). However, the increase in ROS generation was more pronounced upon ALDH7A1 knockdown, either alone or in combination with ALDH3A1 (*p*=0.0001 and 0.0003, respectively), reaffirming its role as an antioxidant under both normoxic and hypoxic conditions. These results suggest that ALDH7A1 upregulation may represent an adaptive response to hypoxia that helps attenuate oxidative stress by limiting ROS accumulation.

Given that ALDH enzyme expression has been correlated with drug resistance to several anticancer drugs [50,51], this study also examined the effect of ALDH1A3, ALDH3A1, and ALDH7A1 knockdown on chemosensitivity in DLD-1 CRC cells. Under normoxic conditions, oxaliplatin (75 µM), irinotecan (75 µM) and 5-FU (100 µM), and treatment resulted in 70%, 85%, and 60% cell kill, respectively, with no significant differences in drug sensitivity between knockdown and control cells. Under hypoxic conditions, cell survival increased from 23% to 88% for oxaliplatin, 21% to 77% for irinotecan and 30% to 67% for 5-FU, indicating increased resistance. Hypoxic cells with ALDH7A1 knockdown showed similar drug responses to controls (p = 0.91, 0.13 and 0.23, for oxaliplatin, irinotecan and 5-FU, respectively), confirming that ALDH7A1 plays no role in drug resistance or hypoxia-mediated reduction in chemosensitivity in CRC cells (Figure 8B).

**Figure 8.**
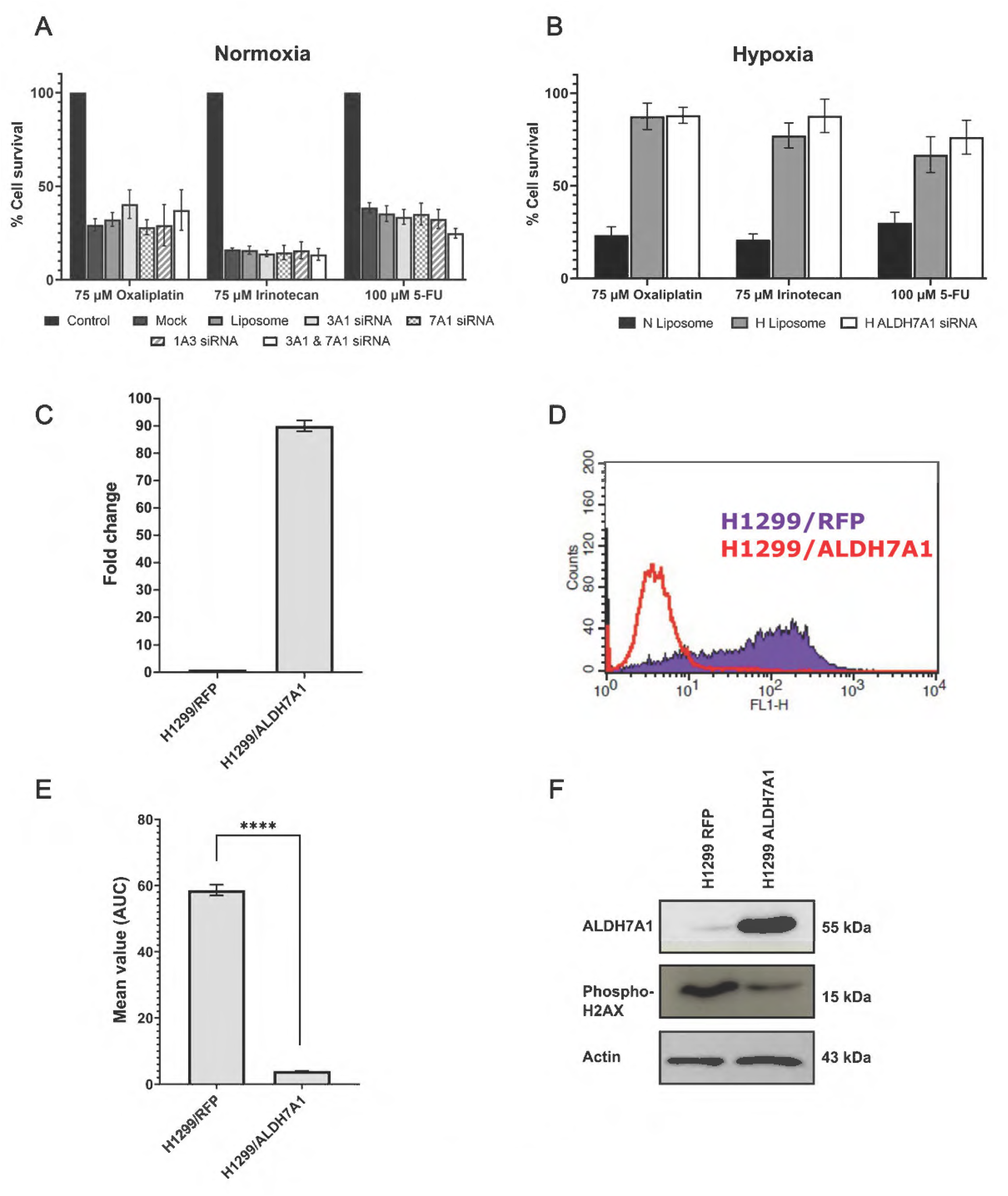
Assessment of chemosensitivity and characterisation of ALDH7A1 overexpression and reactive oxygen species (ROS) production. Cell survival of DLD-1 cells following treatment with oxaliplatin (75 µM), irinotecan (75 µM) and 5-FU (100 µM) for 48 h under normoxic (A) and hypoxic (B) conditions in ALDH isoform siRNA-knockdown cells. Values represent the mean of three independent experiments, and error bars indicate SD. (C) qRT-PCR analysis of ALDH7A1 in H1299/ALDH7A1 cells relative to H1299/RFP controls, expressed as fold change. (D) ROS generation curves in H1299/RFP and H1299/ALDH7A1 cells using FACS, and (E) Geometric mean values of area under the curve (AUC) from ROS production. Values represent the mean of three independent experiments and error bars indicate SD. **** p<0.0001. (F) Western blot analysis of phosphorylated H2AX and ALDH7A1 protein expression in H1299/RFP and H1299/ALDH7A1 cells.

### Overexpression of ALDH7A1 decreases the generation of ROS and prevents DNA damage

To support our observations in the CRC models, an isogenic pair of H1299 non-small cell lung cancer (NSCLC) cell lines with low endogenous ALDH7A1 expression was used, comprising a control and an ALDH7A1-overexpressing cell line [52]. As previously described, H1299 cells were transfected with lentiviral vectors encoding either the full-length cDNA of ALDH7A1 or red fluorescent protein (RFP) to investigate ALDH7A1 in depth. The antioxidant properties of ALDH7A1 were measured in H1299/ALDH7A1 cells using FACS to detect ROS generation. ALDH7A1 overexpression resulted in a significantly reduced formation of ROS compared to the H1299/RFP cells, with more than 90% reduction in ROS generation (Figure 8D and 8E). Expression of phosphorylated H2AX was significantly reduced by approximately 70% in H1299/ALDH7A1 cells compared with H1299/RFP control cells, indicating reduced DNA damage and suggesting that ALDH7A1 may exert a cytoprotective effect (Figure 8F). These collective results further support the conclusion that elevated ALDH7A1 expression in the CRC microenvironment is an adaptive mechanism that promotes its antioxidant role in rescuing cells from intense oxidative stress.

## Discussion

Abnormally high expression of ALDH7A1 has been found in different cancer types, including prostate, ovarian and NSCLC cancers; however, its significance in CRC remains unexplored [40,53,54]. Apart from its vital role in lysine metabolism and osmotic stress, the ALDH7A1 isoform also protects cells from oxidative stress, including lipid peroxidation [55]. The results of this study suggest that among the multiple ALDH isoforms detected in CRC, ALDH7A1 is the most consistently upregulated at both the mRNA and protein levels in response to hypoxia-induced oxidative damage. This was established in multiple settings, including monolayer cell cultures and spheroids, as well as in different CRC cell lines. These findings are consistent with evidence from a recent study by Yang et al., which underscores the role of ALDH7A1 in promoting cellular energy homeostasis by reducing energy consumption under conditions of impaired cellular energy, such as hypoxia and starvation [55]. Energy stress induces the phosphorylation of ALDH7A1 by 5’ AMP-activated protein kinase (AMPK), subsequently delocalising it to the intracellular membrane compartments from the cytosol, leading to broad inhibition of intracellular transport pathways. The NADH generated as a reductive consequence of ALDH7A1 activity inhibits Coat Protein 1 (COP1) vesicle fission by targeting Brefeldin-A ADP-Ribosylated Substrate (BARS), resulting in the abrogation of the COP1 transport [55]. Given the significant role of ALDH7A1 in protecting cells from osmotic, oxidative, and energy stress across diverse pathological conditions, the potential relevance of this isoform in promoting cell survival under hostile conditions such as hypoxia is of interest.

Results from the ALDH7A1 knockdown studies reported here underscore the importance of ALDH7A1 as a key enzyme in combating oxidative stress signals and further emphasise its potential role within the TME of the *in vitro* models employed. The results also suggest that this role may overlap among several ALDH isoforms, primarily ALDH3A1. However, it is important to determine mechanistically whether ALDH7A1 acts independently as an antioxidant enzyme or in concert with other enzymes or pathways in the CRC microenvironment. In addition, given the role of hypoxia and mitochondrial dysfunction in the generation of reactive nitrogen species (RNS) [56], the role of ALDH7A1 in the protection of CRC cells against different types of radicals ought to be considered for evaluation [57]. Collectively, addressing these gaps would broaden understanding of ALDH7A1 and enhance knowledge of how CRC cells adapt to acute or chronic hypoxia.

One of the main challenges in achieving a successful CRC treatment outcome is the presence of intrinsic or acquired drug resistance [58]. It is well known that hypoxia mediates resistance and suppresses the pharmacological activities of both chemotherapy and radiotherapy treatment modalities, which ultimately affects the outcome [59]. This, in part, could be attributed to the ability of hypoxia to enhance the survival of CSCs by acting as a niche that can enhance their dynamic interactions with the microenvironment, promoting metastasis and drug resistance [60]. ALDH1 has been shown to act as a CSC marker in CRC; however, the role of specific ALDH isoforms is not well understood [60]. CSCs might express elevated levels of other ALDH isoforms, such as ALDH7A1, due to their chemoprotective properties and involvement in cell differentiation via retinoic acid pathways, thereby potentially protecting the stem cell component of colon tumours.

Apart from the cytoprotective activity, ALDH7A1 also plays a major role in lysine catabolism in the pipecolic acid pathway, where it catalyses the oxidation of alpha-aminoadipic semialdehyde (α-AASA) to alpha-aminoadipate. A mutation in ALDH7A1 has been linked to pyridoxine-dependent epilepsy (PDE) as a result of defective lysine catabolism [61], which causes accumulating piperidine-6-carboxylate (PC6) to condense with pyridoxal 5’-phosphate (PLP), thereby inactivating this enzyme cofactor that is essential for the normal metabolism of neurotransmitters [61]. As this study’s results suggest that ALDH7A1 expression is an indicator of hypoxia-related changes in early-stage CRC, developing a diagnostic assay to identify patients with these changes may be feasible. The diagnostic kit for PDE patients relies on measuring the α-AASA and PC6 compounds excreted from intracellular pools into urine and plasma, providing a simple way to confirm the diagnosis. In contrast, ALDH7A1 gene analysis enables prenatal diagnosis, as shown in clinical trials [61]. Similarly, such a technology would allow CRC patients after stage I-III surgery to be routinely monitored for signs of tumour recurrence based on hypoxia/ALDH7A1 presence in tissue biopsies and measuring lysine metabolites as surrogate markers in urine and blood samples before and after surgical resection rather than depending on the post-operative serial assays of the carcinoembryonic antigen (CEA) levels which is a widely used tumour marker in CRC patients [62]. Ultimately, earlier detection of aggressive forms of CRC could lead to better treatment options, thereby improving quality of life and survival rates.

In conclusion, the findings demonstrate that ALDH7A1 is upregulated in response to tumour hypoxia and is associated with reduced ROS accumulation, supporting its role as an antioxidant enzyme and a potential mediator of cellular adaptation to hypoxic stress. This further suggests that upregulation of ALDH7A1 under hypoxic conditions may reflect an adaptive response that enhances cancer cell survival and progression. Given the association between ALDH7A1 expression and hypoxia-related changes in early-stage CRC cells, developing screening assays to detect ALDH7A1-associated biomarkers could aid in the early diagnosis of primary colorectal cancer, potentially intercepting disease before it progresses to later stages.

## Acknowledgements

The authors would like to thank Jordan University of Science and Technology and the University of Bradford (RDF & Innovation Pump Priming Scheme) for financial support.

## APPENDIX

**Figure S1.**
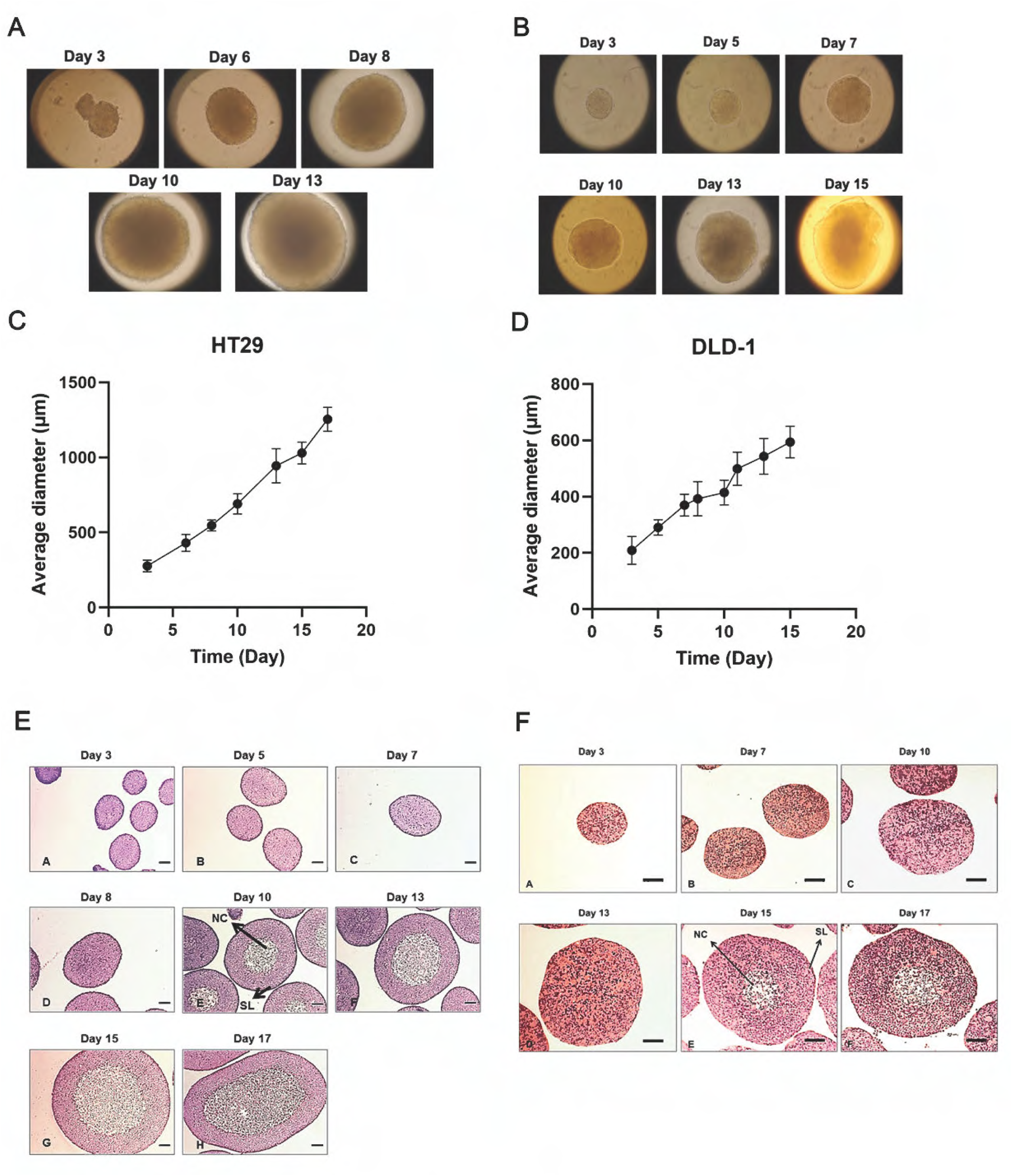
Growth and histology of HT2G (A, C, E) and DLD-1 (B, D, F) spheroids. 3D MCS were generated from HT29 and DLD-1 CRC cell lines, at a concentration of 4 × 10^4^ cells/ml using a spinner flask on a magnetic stirrer plate. (A, B) Photos of HT29 and DLD-1 MCS at x10 magnification, respectively. (C, D) Growth curve of HT29 and DLD-1 MCS, respectively. Points represent the average of at least 20 spheroids, and error bars indicate SD. Spheroids were processed and stained with HCE from Day 3 to Day 17 for HT29 (E) and DLD-1 (F) cell lines. The surface layer (SL) and necrotic core (NC) of MCS were identified. Scale bar = 100µm at 10x objective lens.

**Figure S2.**
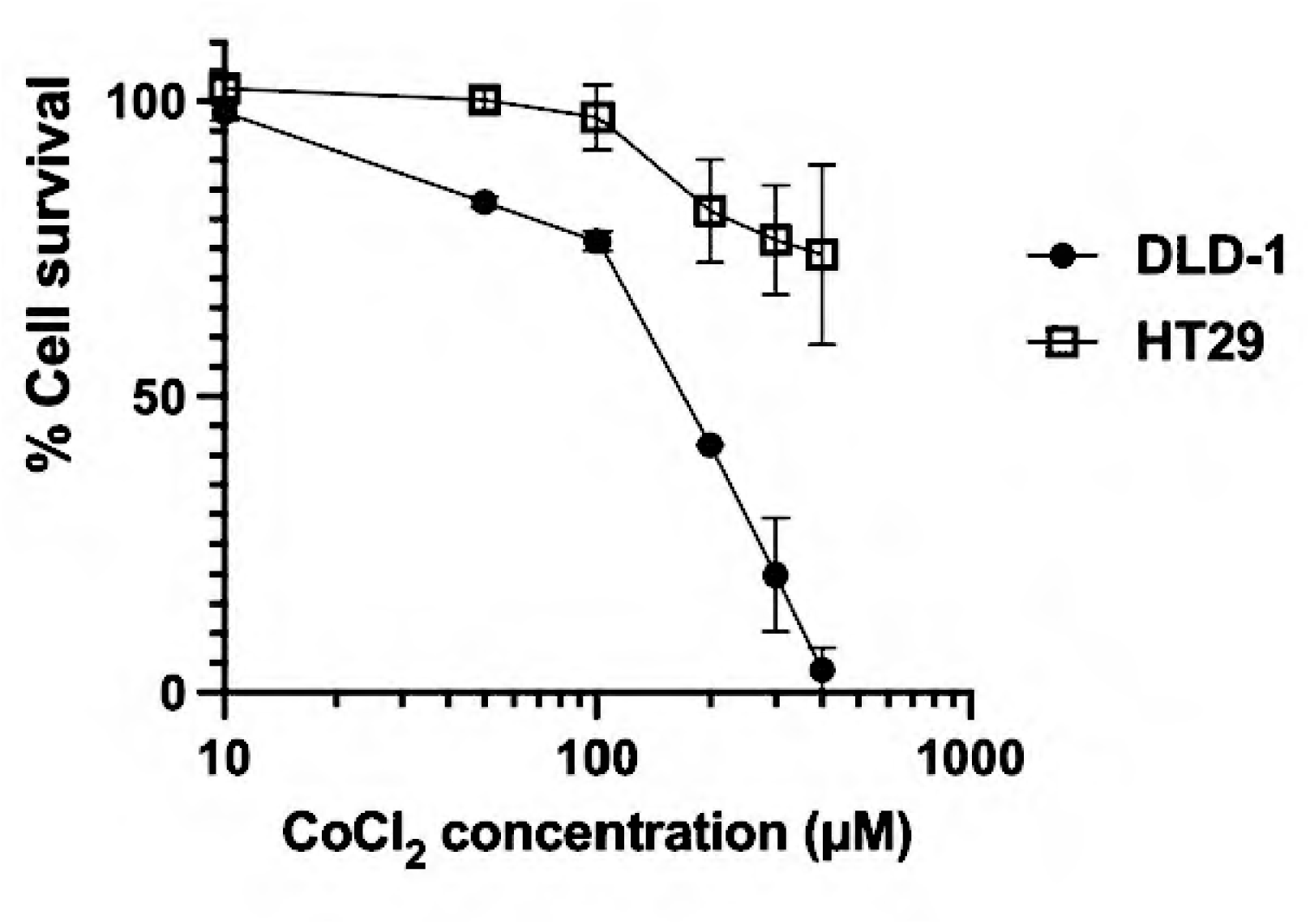
Dose-response curve of CoCl_2_ treatment in HT2G and DLD-1 cell lines. HT29 and DLD-1 cells were seeded and treated with CoCl_2_ (concentration range from 10 µM to 500µM) for 24 h, and percentage cell survival was evaluated using an MTT assay. Values are the mean of three independent experiments, and error bars indicate SD.

**Figure S3.**
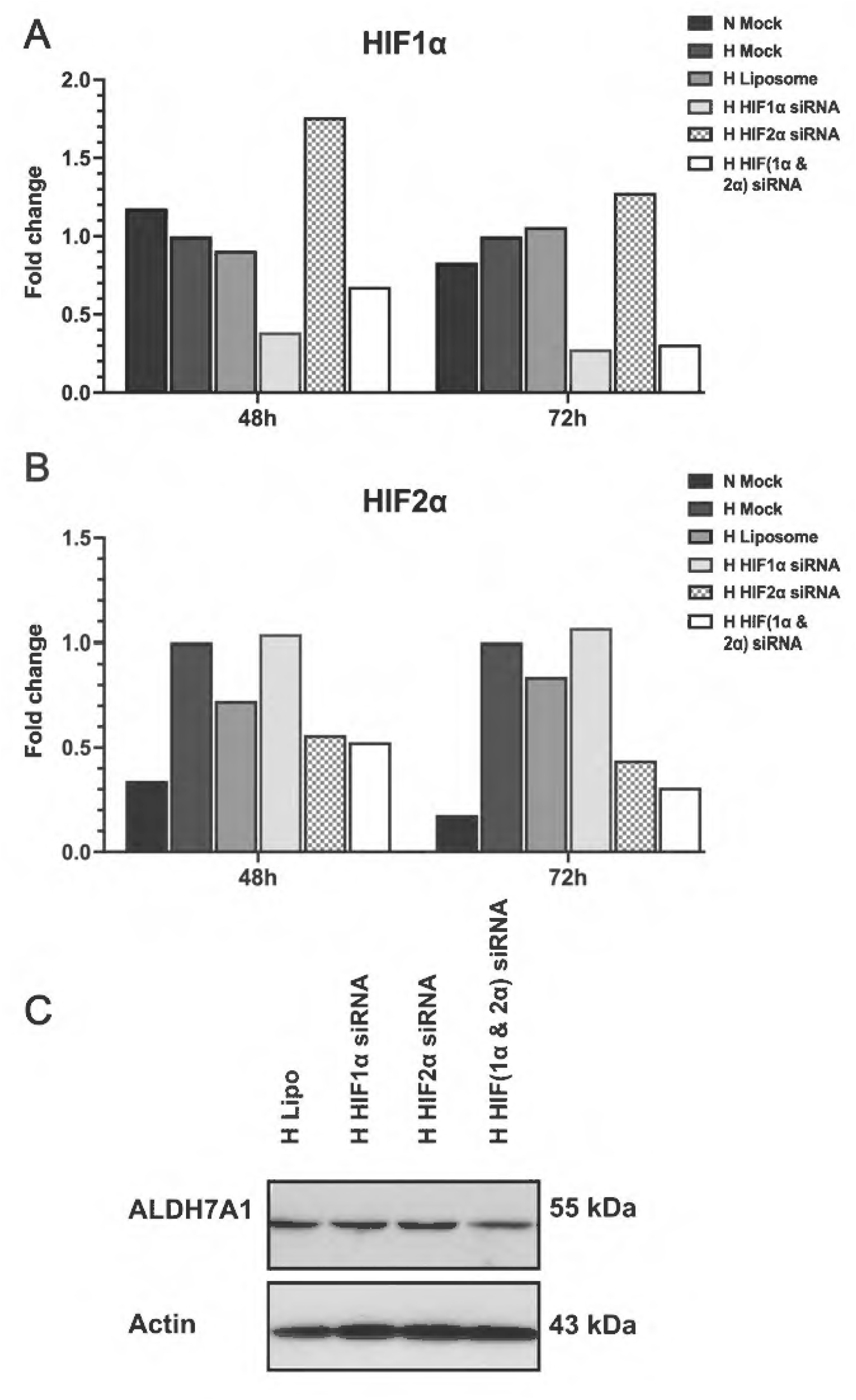
Effect of HIF knockdown on HIF1α and HIF2α mRNA and ALDH7A1 protein expression. HIF1α (A) and HIF2α (B) mRNA expression measured using qRT-PCR after 48 h and 72 h of HIF1α, HIF2α or dual HIF1/2α siRNA-mediated knockdown, under hypoxic conditions. Normoxic mock and hypoxic mock served as basal expression controls under normoxia and hypoxia, respectively. (C) Western blot analysis of ALDH7A1 protein expression after 72 h of HIF knockdown. Actin was used as a loading control. Results represent one experiment.

**Figure S4.**
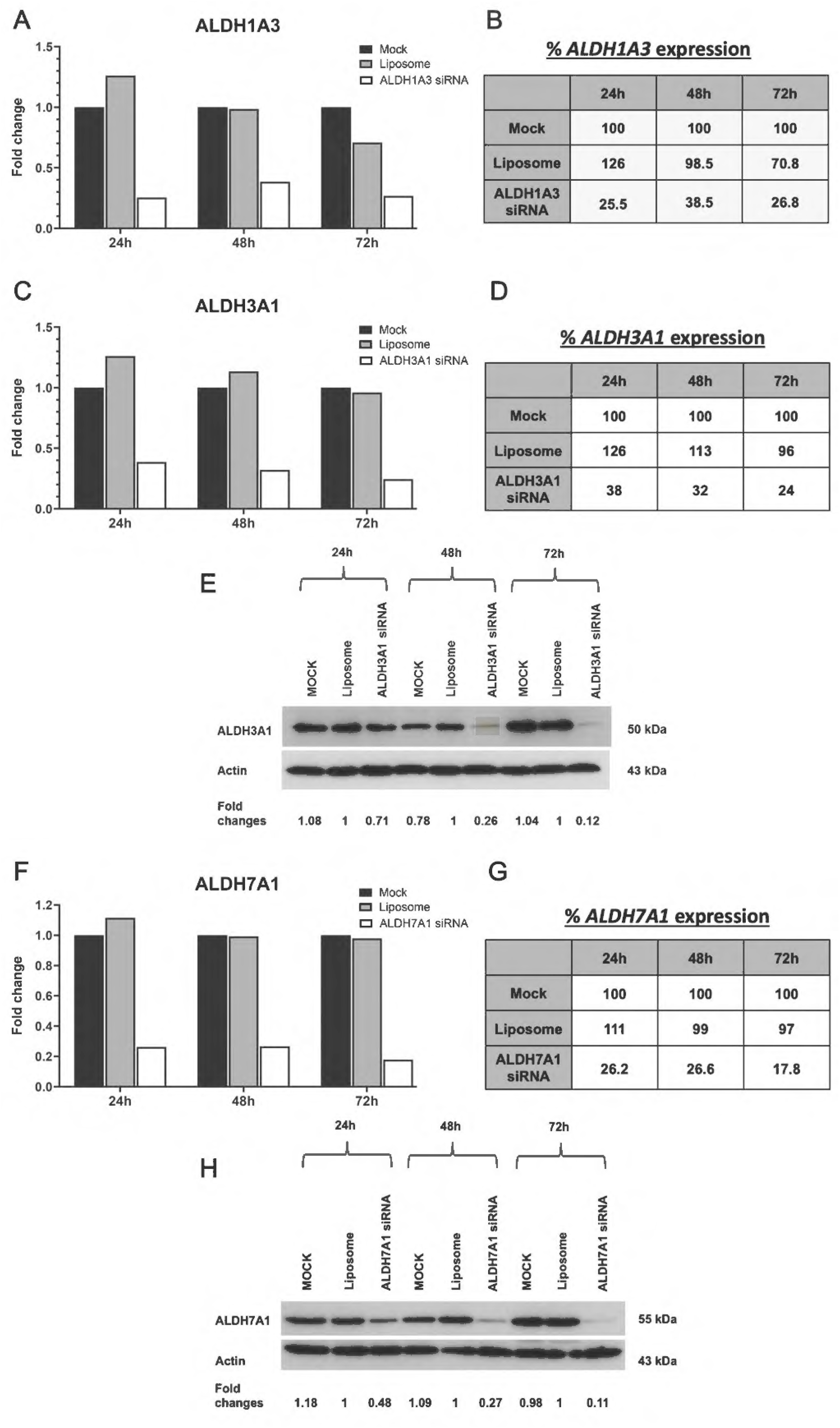
Effect of siRNA knockdown on ALDH mRNA and protein expression in normoxia. DLD-1 cells were transfected with ALDH1A3 (A, B), ALDH3A1 (C-E) and ALDH7A1 (F-H) siRNA under normoxic conditions. mRNA expression was measured using qRT-PCR and protein expression using western blot analysis. (A, C, F) Fold change of ALDH1A3, ALDH3A1, and ALDH7A1 mRNA expression after 24 h, 48 h, and 72 h of transfection with ALDH1A3 siRNA, ALDH3A1 siRNA and ALDH7A1 siRNA, respectively. (B, D, G) Percentage of ALDH1A3, ALDH3A1 and ALDH7A1 mRNA expression, respectively, relative to mock control. (E, H) ALDH3A1 and ALDH7A1 protein expression after 24 h, 48 h and 72 h transfection with ALDH3A1 and ALDH7A1 siRNA, respectively. Fold changes relative to liposome control are indicated below. Actin was used as a loading control.

**Figure S5.**
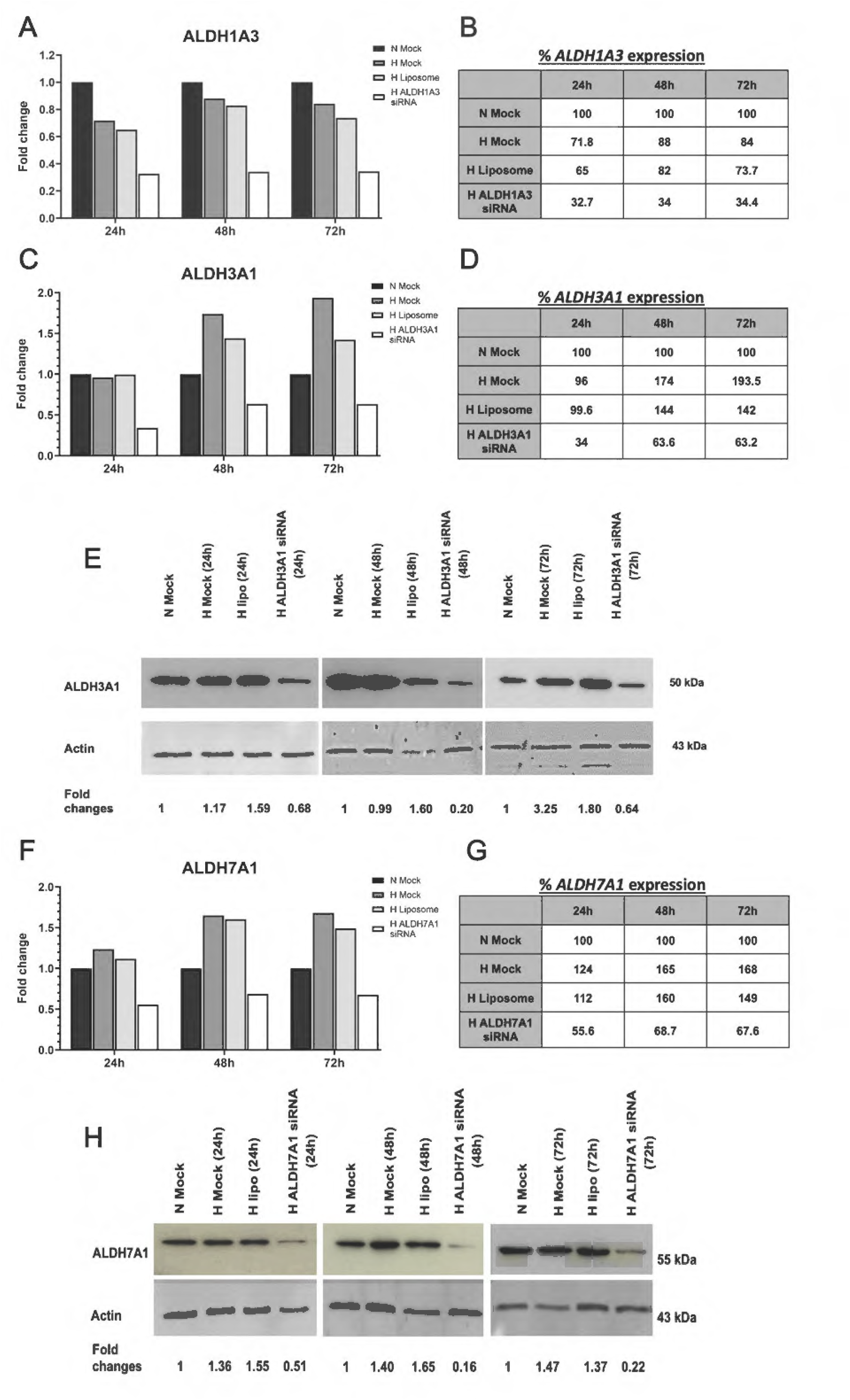
Effect of siRNA knockdown on ALDH mRNA and protein expression in hypoxia. DLD-1 cells were transfected with ALDH1A3 (A, B), ALDH3A1 (C-E) and ALDH7A1 (F-H) siRNA under hypoxic conditions. mRNA expression was measured using qRT-PCR and protein expression using western blot analysis. (A, C, F) Fold change of ALDH1A3, ALDH3A1, and ALDH7A1 mRNA expression after 24 h, 48 h, and 72 h of transfection with ALDH1A3 siRNA, ALDH3A1 siRNA and ALDH7A1 siRNA, respectively, in hypoxia. (B, D, G) Percentage of ALDH1A3, ALDH3A1 and ALDH7A1 mRNA expression, respectively, relative to normoxic mock control. (E, H) ALDH3A1 and ALDH7A1 protein expression after 24 h, 48 h and 72 h transfection with ALDH3A1 and ALDH7A1 siRNA, respectively. Fold changes relative to normoxic mock control indicated below (Mock, Lipo, and siRNA-alone lanes in E and H are shared with Figure 6E and Figure S6G, respectively). Actin used as loading control.

**Figure S6.**
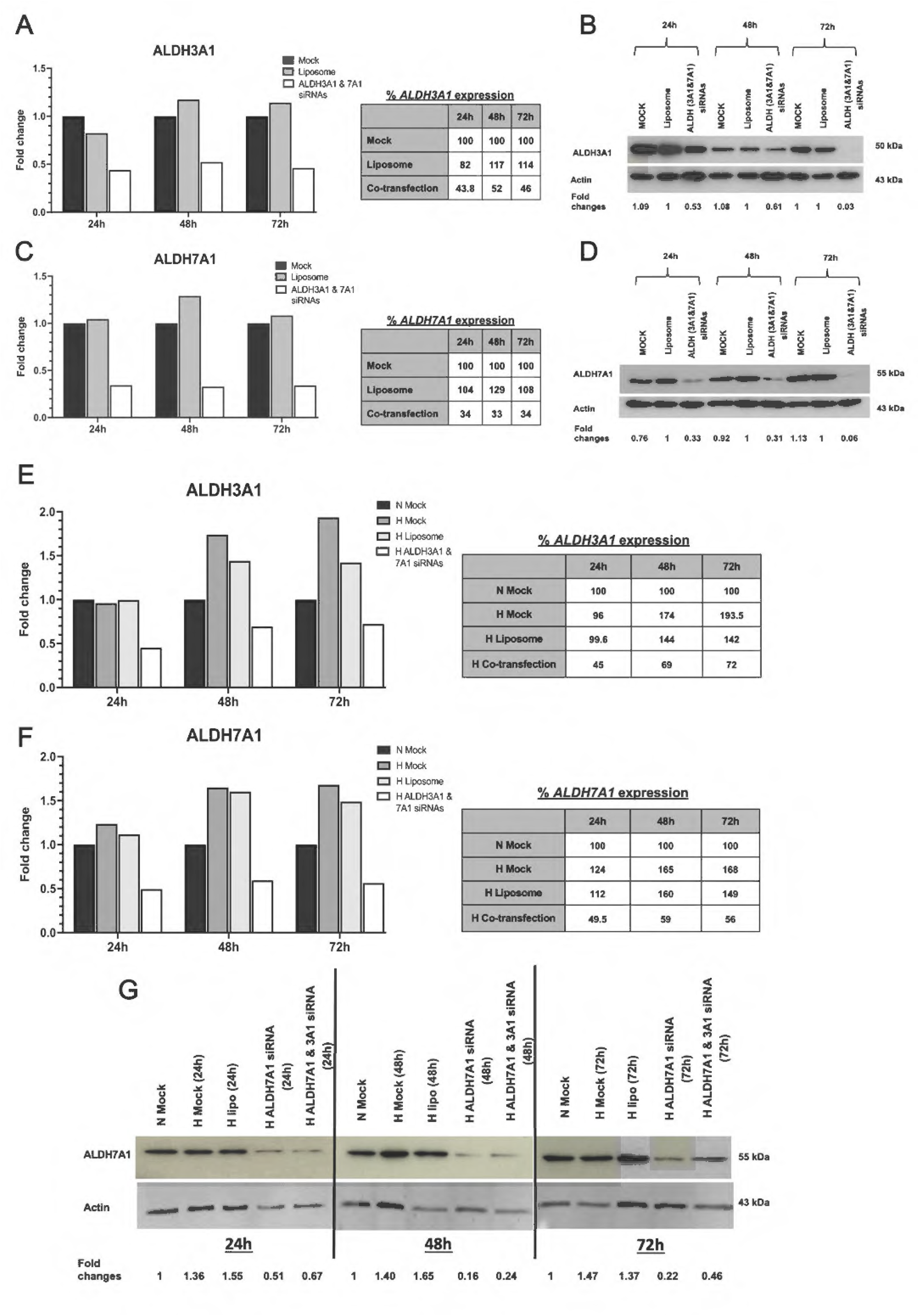
Effect of ALDH3A1/ALDH7A1 siRNA co-transfection on ALDH3A1 and ALDH7A1 expression in DLD-1 cells. DLD-1 cells were co-transfected with ALDH3A1/ALDH7A1 siRNAs for 24 h, 48 h, and 72 h under normoxic (A-D) and hypoxic (E-G) conditions. mRNA and protein expression of each isoform was measured at each time point and condition using qRT-PCR and western blot analysis, respectively. The fold change of ALDH3A1 (A) and ALDH7A1 (C) mRNA expression was measured under normoxic conditions, with the percentage of expression in co-transfected cells relative to mock control indicated in respective tables. ALDH3A1 (B) and ALDH7A1 (D) protein expression in ALDH3A1/7A1 co-transfected cells. Fold changes relative to liposome control are indicated below. Actin was used as a loading control. Fold change of ALDH3A1 (E) and ALDH7A1 (F) expression was measured under hypoxic conditions, with percentage of expression in co-transfected cells relative to normoxic mock control indicated in respective tables. (G) Protein expression of ALDH7A1 in ALDH3A1/7A1 co-transfected cells. Fold changes relative to normoxic mock control indicated below. Actin was used as a loading control.

## References

1. Xi, Y. and Xu, P. (2021) Global colorectal cancer burden in 2020 and projections to 2040. Transl Oncol 14, 101174. 10.1016/j.tranon.2021.101174

2. Ferlay, J. et al. (2021) Cancer statistics for the year 2020: An overview. Int J Cancer. 10.1002/ijc.33588

3. Fink, H. et al. (2026) Global and regional cancer burden attributable to modifiable risk factors to inform prevention. Nat Med 32, 1306–1315. 10.1038/s41591-026-04219-7

4. Haraldsdottir, S. et al. (2014) [Colorectal cancer - review]. Laeknabladid 100, 75–82. 10.17992/lbl.2014.02.531

5. Xie, Y.H. et al. (2020) Comprehensive review of targeted therapy for colorectal cancer. Signal Transduct Target Ther 5, 22. 10.1038/s41392-020-0116-z

6. Khelwatty, S.A. et al. (2025) Advancements in Targeted Therapies for Colorectal Cancer: Overcoming Challenges and Exploring Future Directions. Cancers (Basel) 17. 10.3390/cancers17172810

7. Kawczak, P. and Baczek, T. (2026) Molecular Targeting of EGFR, BRAF, and HER2 Signaling in Colorectal Cancer: Contemporary Advances with Panitumumab, Encorafenib, and Tucatinib. J Clin Med 15. 10.3390/jcm15062387

8. Rohwer, N. et al. (2013) The growing complexity of HIF-1alpha’s role in tumorigenesis: DNA repair and beyond. Oncogene 32, 3569–3576. 10.1038/onc.2012.510

9. Semenza, G.L. (2010) Defining the role of hypoxia-inducible factor 1 in cancer biology and therapeutics. Oncogene 29, 625–634. 10.1038/onc.2009.441

10. Semenza, G.L. (2012) Hypoxia-inducible factors: mediators of cancer progression and targets for cancer therapy. Trends Pharmacol Sci 33, 207–214. 10.1016/j.tips.2012.01.005

11. Kondoh, M. et al. (2013) Hypoxia-Induced Reactive Oxygen Species Cause Chromosomal Abnormalities in Endothelial Cells in the Tumor Microenvironment. PLoS ONE 8, e80349. 10.1371/journal.pone.0080349

12. Fiaschi, T. and Chiarugi, P. (2012) Oxidative stress, tumor microenvironment, and metabolic reprogramming: a diabolic liaison. Int J Cell Biol 2012, 762825. 10.1155/2012/762825

13. Reuter, S. et al. (2010) Oxidative stress, inflammation, and cancer: How are they linked? Free Radical Biology and Medicine 49, 1603–1616. 10.1016/j.freeradbiomed.2010.09.006

14. Balendiran, G.K. et al. (2004) The role of glutathione in cancer. Cell Biochemistry and Function 22, 343–352. 10.1002/cbf.1149

15. Storz, P. (2005) Reactive oxygen species in tumor progression. Front Biosci 10.2741/1667

16. Xanthis, V. et al. (2023) Human Aldehyde Dehydrogenases: A Superfamily of Similar Yet Different Proteins Highly Related to Cancer. Cancers (Basel) 15. 10.3390/cancers15174419

17. Pors, K. and Moreb, J.S. (2014) Aldehyde dehydrogenases in cancer: an opportunity for biomarker and drug development? Drug Discovery Today 19, 1953–1963. 10.1016/j.drudis.2014.09.009

18. Chen, Y. et al. (2011) Aldehyde dehydrogenase 1B1 (ALDH1B1) is a potential biomarker for human colon cancer. Biochemical and Biophysical Research Communications 405, 173–179. 10.1016/j.bbrc.2011.01.002

19. Emmink, B.L. et al. (2013) The secretome of colon cancer stem cells contains drug-metabolizing enzymes. Journal of Proteomics 91, 84–96. 10.1016/j.jprot.2013.06.027

20. Touil, Y. et al. (2014) Colon cancer cells escape 5FU chemotherapy-induced cell death by entering stemness and quiescence associated with the c-Yes/YAP axis. Clin Cancer Res 20, 837–846. 10.1158/1078-0432.CCR-13-1854

21. Tang, X. et al. (2014) A mechanically-induced colon cancer cell population shows increased metastatic potential. Mol Cancer 10.1186/1476-4598-13-131

22. Vasiliou, V. and Nebert, D. (2005) Analysis and update of the human aldehyde dehydrogenase (ALDH) gene family. Human Genomics 2, 138–143

23. Pappa, A. et al. (2003) Aldh3a1 protects human corneal epithelial cells from ultraviolet-and 4-hydroxy-2-nonenal-induced oxidative damage. Free Radical Biology and Medicine 34, 1178–1189. 10.1016/S0891-5849(03)00070-4

24. Brocker, C. et al. (2010) Aldehyde dehydrogenase 7A1 (ALDH7A1) is a novel enzyme involved in cellular defense against hyperosmotic stress. J Biol Chem 285, 18452–18463. 10.1074/jbc.M109.077925

25. Brocker, C. et al. (2011) Aldehyde dehydrogenase 7A1 (ALDH7A1) attenuates reactive aldehyde and oxidative stress induced cytotoxicity. Chem Biol Interact 191, 269–277. 10.1016/j.cbi.2011.02.016

26. Moreb, J.S. et al. (2008) ALDH isozymes downregulation affects cell growth, cell motility and gene expression in lung cancer cells. Mol Cancer 7, 87. 10.1186/1476-4598-7-87

27. O’Connor, K.C. (1999) Three-Dimensional Cultures of Prostatic Cells: Tissue Models for the Development of Novel Anti-Cancer Therapies. Pharmaceutical Research 16, 486–493. 10.1023/a:1011906709680

28. Phillips, R.M. et al. (1994) Increased Activity and Expression of NAD(P)H:Quinone Acceptor Oxidoreductase in Confluent Cell Cultures and within Multicellular Spheroids. Cancer Research 54, 3766–3771

29. Yoshimoto, S. et al. (2023) Establishment of a novel protocol for formalin-fixed paraffin-embedded organoids and spheroids. Biol Open 12. 10.1242/bio.059882

30. Workman, P. et al. (2010) Guidelines for the welfare and use of animals in cancer research. Br J Cancer 102, 1555–1577. 10.1038/sj.bjc.6605642

31. Law, P.C. et al. (2012) Astragalus saponins downregulate vascular endothelial growth factor under cobalt chloride-stimulated hypoxia in colon cancer cells. BMC Complement Altern Med 12, 160. 10.1186/1472-6882-12-160

32. Sieuwerts, A.M. et al. (1995) The MTT tetrazolium salt assay scrutinized: how to use this assay reliably to measure metabolic activity of cell cultures in vitro for the assessment of growth characteristics, IC50-values and cell survival. Eur J Clin Chem Clin Biochem 33, 813–823. 10.1515/cclm.1995.33.11.813

33. Allison, S.J. and Milner, J. (2014) RNA Interference by Single-and Double-stranded siRNA With a DNA Extension Containing a 3’ Nuclease-resistant Mini-hairpin Structure. Mol Ther Nucleic Acids 2, e141. 10.1038/mtna.2013.68

34. Soehngen, E. et al. (2014) Hypoxia upregulates aldehyde dehydrogenase isoform 1 (ALDH1) expression and induces functional stem cell characteristics in human glioblastoma cells. Brain Tumor Pathol 31, 247–256. 10.1007/s10014-013-0170-0

35. Huang, J. et al. (2024) ALDH1A3 contributes to tumorigenesis in high-grade serous ovarian cancer by epigenetic modification. Cell Signal 116, 111044. 10.1016/j.cellsig.2024.111044

36. Hirschhaeuser, F. et al. (2010) Multicellular tumor spheroids: an underestimated tool is catching up again. J Biotechnol 148, 3–15. 10.1016/j.jbiotec.2010.01.012

37. Sutherland, R.M. (1998) Tumor hypoxia and gene expression--implications for malignant progression and therapy. Acta Oncol 37, 567–574. 10.1080/028418698430278

38. Imamura, T. et al. (2009) HIF-1alpha and HIF-2alpha have divergent roles in colon cancer. Int J Cancer 124, 763–771. 10.1002/ijc.24032

39. Piret, J.P. et al. (2002) CoCl2, a chemical inducer of hypoxia-inducible factor-1, and hypoxia reduce apoptotic cell death in hepatoma cell line HepG2. Ann N Y Acad Sci 973, 443–447. 10.1111/j.1749-6632.2002.tb04680.x

40. van den Hoogen, C., et al. (2011) The aldehyde dehydrogenase enzyme 7A1 is functionally involved in prostate cancer bone metastasis. Clin Exp Metastasis 28, 615–625. 10.1007/s10585-011-9395-7

41. Goda, N. et al. (2003) HIF-1 in cell cycle regulation, apoptosis, and tumor progression. Antioxid Redox Signal 5, 467–473. 10.1089/152308603768295212

42. Marchitti, S.A. et al. (2011) Ultraviolet radiation: cellular antioxidant response and the role of ocular aldehyde dehydrogenase enzymes. Eye Contact Lens 37, 206–213. 10.1097/ICL.0b013e3182212642

43. Klaunig, J.E. et al. (2010) Oxidative stress and oxidative damage in carcinogenesis. Toxicol Pathol 38, 96–109. 10.1177/0192623309356453

44. Dando, I. et al. (2015) Antioxidant Mechanisms and ROS-Related MicroRNAs in Cancer Stem Cells. Oxid Med Cell Longev 2015, 425708. 10.1155/2015/425708

45. Lassen, N. et al. (2007) Multiple and additive functions of ALDH3A1 and ALDH1A1: cataract phenotype and ocular oxidative damage in Aldh3a1(-/-)/Aldh1a1(-/-) knock-out mice. J Biol Chem 282, 25668–25676. 10.1074/jbc.M702076200

46. Li, Y. et al. (1994) DNA damage caused by reactive oxygen species originating from a copper-dependent oxidation of the 2-hydroxy catechol of estradiol. Carcinogenesis 15, 1421–1427. 10.1093/carcin/15.7.1421

47. Kuo, L.J. and Yang, L.X. (2008) Gamma-H2AX - a novel biomarker for DNA double-strand breaks. In Vivo 22, 305–309

48. Liu, Q. et al. (2004) A Fenton reaction at the endoplasmic reticulum is involved in the redox control of hypoxia-inducible gene expression. Proc Natl Acad Sci U S A 101, 4302–4307. 10.1073/pnas.0400265101

49. Lopez-Barneo, J. et al. (2001) Cellular mechanism of oxygen sensing. Annu Rev Physiol 63, 259–287. 10.1146/annurev.physiol.63.1.259

50. Su, Y. et al. (2010) Aldehyde dehydrogenase 1 A1-positive cell population is enriched in tumor-initiating cells and associated with progression of bladder cancer. Cancer Epidemiol Biomarkers Prev 19, 327–337. 10.1158/1055-9965.EPI-09-0865

51. Januchowski, R. et al. (2013) The role of aldehyde dehydrogenase (ALDH) in cancer drug resistance. Biomed Pharmacother 67, 669–680. 10.1016/j.biopha.2013.04.005

52. Moreb, J.S. et al. (2012) The enzymatic activity of human aldehyde dehydrogenases 1A2 and 2 (ALDH1A2 and ALDH2) is detected by Aldefluor, inhibited by diethylaminobenzaldehyde and has significant effects on cell proliferation and drug resistance. Chem Biol Interact 195, 52–60. 10.1016/j.cbi.2011.10.007

53. van den Hoogen, C., et al. (2010) High aldehyde dehydrogenase activity identifies tumor-initiating and metastasis-initiating cells in human prostate cancer. Cancer Res 70, 5163–5173. 10.1158/0008-5472.CAN-09-3806

54. Saw, Y.T. et al. (2012) Characterization of aldehyde dehydrogenase isozymes in ovarian cancer tissues and sphere cultures. BMC Cancer 12, 329. 10.1186/1471-2407-12-329

55. Yang, J.S. et al. (2019) ALDH7A1 inhibits the intracellular transport pathways during hypoxia and starvation to promote cellular energy homeostasis. Nat Commun 10, 4068. 10.1038/s41467-019-11932-0

56. Reuter, S. et al. (2010) Oxidative stress, inflammation, and cancer: how are they linked? Free Radic Biol Med 49, 1603–1616. 10.1016/j.freeradbiomed.2010.09.006

57. Haklar, G. et al. (2001) Different kinds of reactive oxygen and nitrogen species were detected in colon and breast tumors. Cancer Lett 165, 219–224. 10.1016/s0304-3835(01)00421-9

58. Holohan, C. et al. (2013) Cancer drug resistance: an evolving paradigm. Nat Rev Cancer 13, 714–726. 10.1038/nrc3599

59. Wouters, A. et al. (2007) Review: implications of in vitro research on the effect of radiotherapy and chemotherapy under hypoxic conditions. Oncologist 12, 690–712. 10.1634/theoncologist.12-6-690

60. Heddleston, J.M. et al. (2010) Hypoxia inducible factors in cancer stem cells. Br J Cancer 102, 789–795. 10.1038/sj.bjc.6605551

61. Scharer, G. et al. (2010) The genotypic and phenotypic spectrum of pyridoxine-dependent epilepsy due to mutations in ALDH7A1. J Inherit Metab Dis 33, 571–581. 10.1007/s10545-010-9187-2

62. Walker, A.S. et al. (2014) Future directions for the early detection of colorectal cancer recurrence. J Cancer 5, 272–280. 10.7150/jca.8871

